# ARHGEF7 S-glutathionylation promotes cancer cell migration through Rac1 activation

**DOI:** 10.64898/2026.05.01.722049

**Authors:** William H. Schiff, Madhu C. Shivamadhu, Faezeh Mashhadi Ramezani, Dhanushika S. K. Kukulage, Rayavarapu Padmavathi, Young-Hoon Ahn

**Affiliations:** Department of Chemistry, Drexel University, Philadelphia, PA 19104, USA

**Keywords:** ARHGEF7, β-PIX, S-glutathionylation, redox signaling, reactive oxygen species, migration, invasion

## Abstract

Reactive oxygen species (ROS) are central signaling molecules in many biological processes by inducing oxidative modifications of protein cysteine residues, including S-glutathionylation. Increasing evidence supports that ROS contribute to cancer progression via promoting cancer cell migration, invasion, and metastasis. Nevertheless, the protein targets of S-glutathionylation that regulate cancer cell motility remain ill-defined. In this study, we report on the redox regulation of ARHGEF7, a guanine nucleotide exchange factor highly expressed in metastatic cancer cells, that plays a major role in regulating cell migration. Our data demonstrates that ARHGEF7 is selectively glutathionylated at the highly conserved C312 residue in its PH domain, which is implicated in regulating its enzymatic activity. Breast cancer cell lines showed increased cell migration and invasion upon glutathionylation of ARHGEF7 at C312 in response to both oxidative stress and epidermal growth factor (EGF). Mechanistically, upon C312 glutathionylation, ARHGEF7 exhibited significantly enhanced binding to Rac1 and increased Rac1 recruitment to the cell membrane and lamellipodia. ARHGEF7 S-glutathionylation also increased its enzymatic rate of GDP-GTP nucleotide exchange, resulting in Rac1 activation. Consequently, ARHGEF7 C312 S-glutathionylation induced Rac1-PAK1 activation and their downstream pathways, including LIMK1 and MEK1, thereby enhancing migration and invasion. Our data reveal a new redox player in cell migration, with its potential implications for ROS-induced cancer progression.

## Introduction

Cell migration is a fundamental, controlled mechanism essential for many physiological processes, including embryonic development, immune responses, and wound healing, and is critical for tissue formation and maintenance.^1,2^ In contrast, aberrations in cell migration contribute to pathological processes, including cancer metastasis and chronic inflammation.^3-5^ Accumulating evidence indicates that cell migration, as well as its associated processes such as invasion and cell-cell adhesion, are regulated by redox signaling, especially with reactive oxygen species acting as signaling molecules.^6,7^ For example, H_2_O_2_ is generated at the site of injury as a signal to recruit leukocytes during wound healing.^8^ H_2_O_2_ is locally produced at the leading edge of migrating cells, inducing protrusion formation.^9^ ROS regulate multiple signaling pathways implicated in cell migration and invasion, including mitogen-activated protein kinases (MAPKs),^10^ focal adhesion complexes,^11^ and Rho GTPases (e.g., Rac1 and RhoA),^12^ which control cytoskeletal remodeling and cell-cell adhesion to enhance cell migration.^7^ Extensive evidence indicates that ROS, oxidative stress, or oxidase expression induces migration in various cell types, including epithelial, fibroblast, and endothelial cells, and promotes the progression and metastasis of cancer cells.^13-21^

At a molecular level, ROS regulate biological processes in part via inducing protein cysteine oxidation and oxidative modifications, including sulfenylation, intramolecular disulfide formation, and S-glutathionylation.^22^ As one of the primary cysteine modifications, protein S-glutathionylation has been demonstrated to regulate major biological processes,^23-26^ including cell migration, invasion, and epithelial-mesenchymal transition (EMT). For example, previous studies have reported that S-glutathionylation of actin and the calcium-binding protein S100A4 regulates cell migration by modulating actin and myosin filament formation in cancer cells.^27,28^ Glutathionylation of Ras leads to its activation, potentially modulating cell proliferation and migration.^29^ In addition, S-glutathionylation of 15-PGDH inhibits its activity, which is associated with increasing EMT.^30^ We reported that S-glutathionylation of the Ser/Thr phosphatase PP2Cα,^31^ the cytoskeletal adaptor protein NISCH,^32^ the fatty acid-binding protein FABP5,^33^ and E-cadherin regulator p120-catenin^34^ regulates mitogenic signaling, cytoskeletal structures, and cell-cell adhesion, thereby increasing the migration and invasion of epithelial cancer cells. Nevertheless, specific proteins that regulate cell migration and invasion via S-glutathionylation, particularly in metastatic cancer cells, remain largely elusive.

An early seminal observation in redox signaling is that growth factors, such as PDGF, produce intracellular hydrogen peroxide (H_2_O_2_),^35^ which boosts kinase signaling by inactivating protein-tyrosine phosphatases (PTPs).^36,37^ Consistently, epidermal growth factor receptor (EGFR), an important regulator of cell migration and proliferation, has been reported for its roles in redox signaling.^38,39^ Specifically, EGFR activates the Ras superfamily, a group of small GTPases that regulate cell migration.^40,41^ The Rho subfamily of the Ras superfamily (e.g., Rac1, Cdc42, and RhoA) regulates cell adhesion and motility through reorganization of the actin cytoskeleton and stress fibers.^42^ Importantly, EGF activates Rac1 GTPase, which promotes cell migration by forming and stabilizing lamellipodia and membrane ruffles.^43-47^ Active Rac1 is an essential subunit that binds to and activates NADPH oxidases, which elevate intracellular ROS levels, thereby initiating EGF-induced redox signaling.^48-51^ In addition, EGFR is activated by ROS-induced cysteine oxidation, propagating its downstream signaling.^52^ Therefore, the EGFR-Rac1-ROS axis creates a positive feed-forward loop that amplifies redox signaling and enhances cell migration and invasion.^53^

In this report, we demonstrate glutathionylation of the Rho guanine nucleotide exchange factor 7 (ARHGEF7, also known as PAK-interacting exchange factor beta, β-PIX), which plays a primary role in promoting cell migration and invasion via activating Rac1.^54^ The GTPases, including Rac1, are regulated by two competing classes of enzymes: GTPase-activating proteins (GAPs) that accelerate GTP hydrolysis to turn off signaling, and guanine nucleotide exchange factors (GEFs) that facilitate GDP-GTP exchange to activate signaling.^55^ ARHGEF7 is a member of the Dbl family of GEFs, characterized by containing tandem Dbl-homology (DH) and pleckstrin-homology (PH) domains, and catalyzes the GDP-GTP exchange for activation of Rac1.^54,56^ During cell migration, ARHGEF7, in complex with GIT1, is recruited to integrin-associated focal complexes at the cell protrusion, where it binds and activates Rac1.^57-59^ The active Rac1 stimulates lamellipodial protrusions via activating the Arp2/3 and cortactin-mediated actin branching.^60^ Rac1 also activates its effector protein, PAK1, and its downstream LIMK1-cofilin axis, thereby stimulating actin filament formation.^60,61^ PAK1 also activates the MEK1-ERK1/2 pathway, promoting cell survival, growth, and migration.^62^ Consistent with its role in activating two established oncogenes (i.e., Rac1 and PAK1),^63^ ARHGEF7 is expressed at high levels in metastatic cancers, including breast and colorectal cancers, as well as renal cell carcinoma.^64-67^ Dysregulation of the ARHGEF7-Rac1-PAK1 complex has been noted with increased cancer aggressiveness correlated with both increased Rac1^68,69^ and ARHGEF7 activities.^67,70^

In this report, our data demonstrate that ARHGEF7 is selectively glutathionylated at cysteine 312 (C312) in the PH domain in response to EGF or oxidative stress, a multi-modal domain implicated in binding to phosphoinositide or regulating its GEF enzymatic activity.^71^ ARHGEF7 C312 glutathionylation increases the migration and invasion of MDA-MB-231 breast cancer cells. Mechanistically, we demonstrate that ARHGEF7 C312 glutathionylation confers two effects: first, ARHGEF7 glutathionylation enhances ARHGEF7’s interactions with Rac1, thereby recruiting Rac1 to cell membranes and protrusions. Second, ARHGEF7 glutathionylation increases its enzymatic activity, resulting in the activation of Rac1. Our biochemical analysis suggests potential conformational changes of ARHGEF7 upon S-glutathionylation that enhance Rac1 binding and activation, thereby increasing cell migration.

## Results

### ARHGEF7 is susceptible to glutathionylation at conserved C312

Previously, we sought to identify proteins that regulate cell migration via S-glutathionylation.^31^ To do so, we analyzed a proteomic database of S-glutathionylation, selected candidate proteins using bioinformatics, and performed cell migration assays, which identified proteins, including ARHGEF7, that increase cell migration under oxidative stress (i.e., limited glucose).^31^ Specifically, our proteomics identified two glutathionylated cysteine sites in ARHGEF7, i.e., C312 and C543 (Figure S1). Our cell migration analysis demonstrated that ARHGEF7 wild-type (WT) increases MDA-MB-231 cell migration under oxidative stress, whereas its cysteine mutant (C312S/C543S) abolishes the increase, suggesting a potential role of ARHGEF7 cysteine oxidation in cell migration.^31^

ARHGEF7 (i.e., β-PIX) comprises multiple domains (Figure 1A and S2A):^54^ the N-terminal calponin homology (CH) domain mediates protein-protein interactions that control ARHGEF7 localization. The SH3 domain binds and recruits Rac1 and PAK1 to protrusion.^57,72^ The DH and PH domains are responsible for GEF enzymatic activity.^56,73,74^ The PH domain is also implicated in binding to phosphoinositide, e.g., PI(3,5)P_2_, which may target ARHGEF7 to specific compartments.^75,76^ The GIT-binding domain (GBD), composed of an α-helix, binds GIT, a GAP that interacts with paxillin, a central participant in the focal adhesion complex (comprising Talin, vinculin, and integrin) at the focal adhesion (see Figure 5).^77^ The C-terminus is largely disordered but contains a coiled-coil (CC) domain that mediates ARHGEF7 oligomerization (i.e., trimers).^78^ Along with dimeric GIT, trimeric ARHGEF7 forms a large complex in focal adhesion or condensates.^79^ Interestingly, ARHGEF7 has multiple isoforms (at least 8, designated isoform “a-f” in the NCBI gene) (Figure S2A-B). All isoforms contain identical SH3-DH-PH-GBD domains but vary in C-terminal sequences or lack the N-terminal CH3 domain (Figure S2A-B). The canonical ARHGEF7 (human isoform c) contains the CH3 domain and a disordered C-terminus without the CC domain (Figure S2B). In contrast, the most ubiquitous form of ARHGEF7 (isoform a) lacks CH3 but contains a disordered C-terminus with a CC domain (Figure S2B). Two cysteines identified for S-glutathionylation are found in the PH (i.e., C312) and GBD (i.e., C543) domains (numbering in isoform-a, corresponding to C490 and C721, respectively, in canonical isoform-c). C312 is positioned at the β1-β2 loop in the PH domain (Figure 1A), surrounded by basic residues, which are predicted to interact with phosphoinositide.^75^ On the other hand, C543 is located within the α-helical GIT-binding domain (aa 517-550) and is in proximity to the GIT binding interface (Figure S2C). ARHGEF7 contains 10 cysteines across its domains (human isoform a, out of 646 amino acids), with 7 cysteines in the DH-PH domains. Both C312 and C543 are predicted to have accessible surface area (ASA) and pK_a_ values comparable to other cysteines (ASA = 77% and 100%; pK_a_ = 9.7 and 9.3 for C312 and C543, respectively, Figure 1B and S2D). Both C312 and C543 are conserved among mammals, with little or no differences in the surrounding residues (Figure 1C and S3A). These two cysteines are conserved or closely aligned with cysteines in the related family enzyme, e.g., ARHGEF6 (also known as α-PIX) (Figure S3B-C).

**Figure 1.**
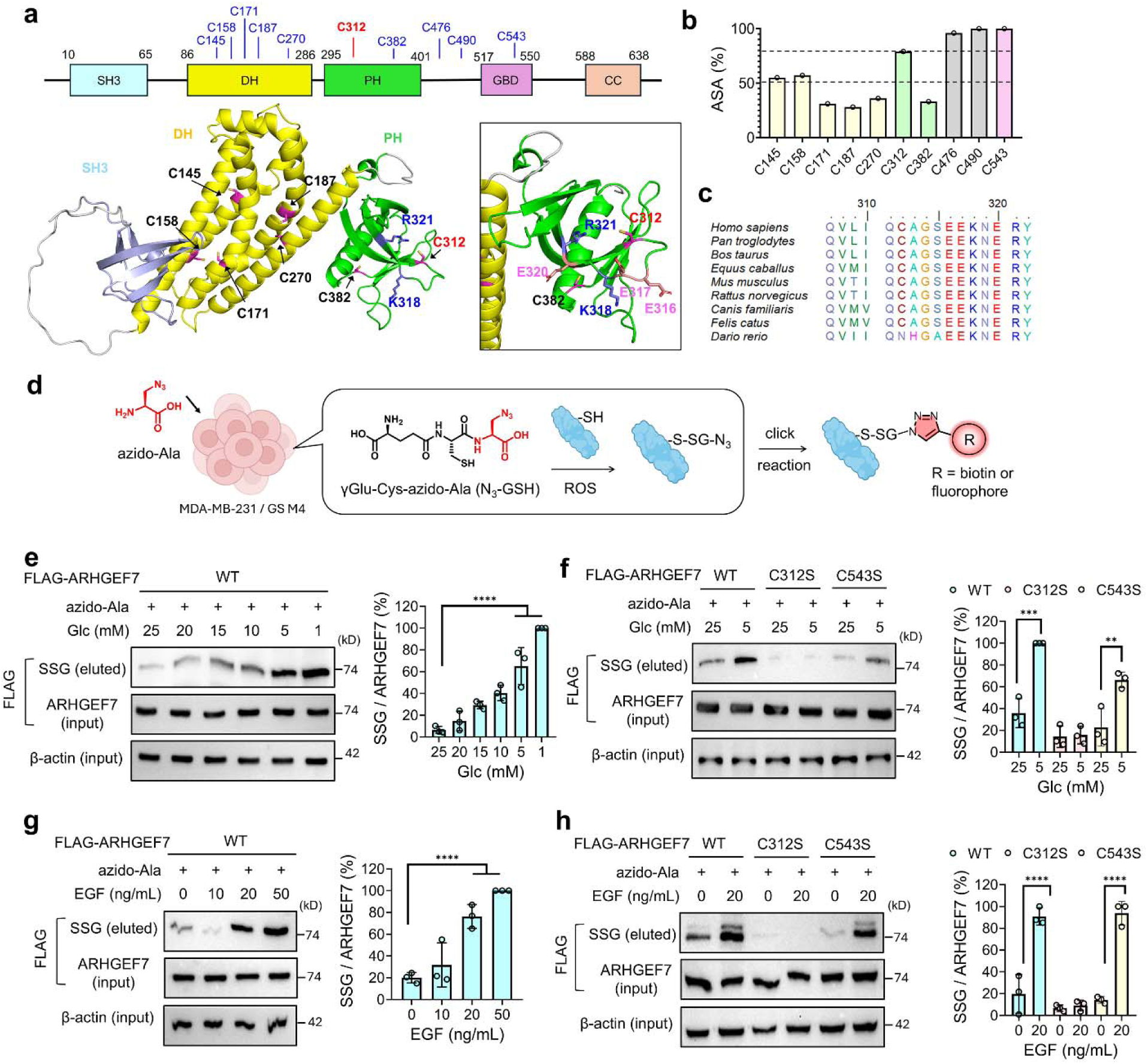
ARHGEF7 is selectively glutathionylated at the conserved C312. (**a**) ARHGEF7 (human isoform-a, 646 residues, NM_001113513) domains, structure, and selected residues. The structure is an AlphaFold model (AF-Q14155-1-F1-model_v6_isoform_a) of ARHGEF7 showing the structured SH3, DH, and PH domains with cysteines and nearby residues. (**b**) Accessible surface areas (ASA) of cysteines in ARHGEF7. (**c**) Sequence alignments around C312 in ARHGEF7 orthologs. (**d**) Clickable glutathione approach. Cells expressing a glutathione synthetase mutant (GS M4) synthesize and produce clickable glutathione (N_3_-GSH) using endogenous γGlu-Cys and exogenous azido-Ala. Clickable glutathione forms S-glutathionylation with proteins, which are analyzed after click reactions. (**e**-**f**) Glutathionylation of ARHGEF7 WT and cysteine mutants in MDA-MB-231 cells in response to high glucose (HG, 25 mM) or low glucose (LG, 5 mM) conditions: cells were incubated for 20 h (n=3). (**g**-**h**) Glutathionylation of ARHGEF7 constructs in MDA-MB-231 cells upon adding EGF (n=3). EGF was incubated for 20 h. The lysates were used for click reactions with biotin-alkyne and analyzed by western blot with FLAG antibody before (input) and after (eluted) pull-down with streptavidin-agarose. Data represent the mean ± SD. The statistical difference was analyzed by one-way ANOVA and Tukey’s post-hoc test (**e**-**h**), where *p < 0.03, **p < 0.002, ***p < 0.0002, ****p < 0.0001.

We analyzed ARHGEF7 glutathionylation in MDA-MB-231 cells, especially using our clickable glutathione approach.^80^ In this approach, clickable glutathione (azido-glutathione, N_3_-GSH) is synthesized in cells expressing a glutathione synthetase mutant (GS M4) upon incubation with azido-Ala. When clickable glutathione forms S-glutathionylation, glutathionylated proteins are analyzed after click reaction with biotin- or fluorophore-alkyne (Figure 1D).^81^ Following application of GS M4 and azido-Ala, MDA-MB-231 cells were exposed to oxidative stress (i.e., limited glucose). Previously, we have shown that low or no glucose concentrations (0-5 mM) increase ROS and induce global S-glutathionylation in various cell lines grown and maintained in high-glucose conditions (i.e., 25 mM glucose).^31,32,82,83^ Limited glucose availability likely reduces NADPH biosynthesis, thereby preventing the maintenance of redox homeostasis in glucose-addicted cancer cells.^84,85^ ARHGEF7 increased its glutathionylation as the glucose gradient decreased (Figure 1E). To determine specific cysteine sites of S-glutathionylation, MDA-MB-231 cells were transfected with either WT or a single cysteine mutant (C312S or C543S) of ARHGEF7. Under low glucose conditions, ARHGEF7 WT was observed to be glutathionylated. ARHGEF7 C543S also increased glutathionylation, albeit at apparently slightly lower levels than WT. In contrast, ARHGEF7 C312S displayed little to no induction of glutathionylation (Figure 1F). To increase the relevance of S-glutathionylation to cell migration, we monitored ARHGEF7 glutathionylation upon incubating EGF for a long duration (20 h). ARHGEF7 showed a dose-dependent increase of S-glutathionylation (Figure 1G). Importantly, ARHGEF7 WT and C543S displayed strong levels of S-glutathionylation, whereas no significant glutathionylation was observed with C312S (Figure 1H). The same results were observed for ARHGEF7 glutathionylation in the presence of EGF incubated only for a short duration (30 min) (Figure S4A).

To corroborate the data, ARHGEF7 glutathionylation was examined *in vitro* using purified ARHGEF7 constructs comprising the DH-PH domains (ARHGEF7 DP) or the SH3-DH-PH domains (ARHGEF7 SDP). Both WT and C312S versions of ARHGEF7 constructs (DP and SDP) were incubated with hydrogen peroxide (H_2_O_2_). Both DP and SDP constructs showed increasing glutathionylation in WT, whereas showing little induction of glutathionylation in C312S by the H_2_O_2_ gradient (Figure S4B-C). Similarly, oxidized glutathione (GSSG) induced a significant modification in the ARHGEF7 SDP WT construct, compared to C312S (Figure S4D). These data support the conclusion that ARHGEF7 C312 is selectively susceptible to glutathionylation under physiological growth-factor stimulation and oxidative-stress conditions.

### ARHGEF7 glutathionylation increases cell migration and invasion

Next, we investigated the effect of ARHGEF7 glutathionylation on cell migration and invasion. MDA-MB-231 cells expressing ARHGEF7 WT, C312S, or C543S were grown and incubated under either high-glucose (25 mM, HG) or low-glucose (5 mM, LG) conditions and assessed for cell migration using a wound-healing assay (Figure 2A). MDA-MB-231 cells expressing ARHGEF7 WT showed a significant increase in cell migration in LG versus HG (bar 4 vs 3). Similar levels of increase in cell migration were also observed in cells expressing C543S (bar 8 vs. 7). However, cells expressing ARHGEF7 C312S did not show increases as significant as WT and C543S (bars 6 vs. 5), while exhibiting comparable increases to cells without ectopic expression of ARHGEF7 (bars 2 vs. 1), suggesting that glutathionylation at C312 is accountable for the increase in cell migration. Under these conditions (i.e., LG vs. HG for 24 h), cell numbers or viability did not change significantly (Figure S5A-B), suggesting that cell proliferation was unaffected or minimally affected. In addition to limited glucose conditions, MDA-MB-231 cells were stimulated with EGF, showing similar results: both ARHGEF7 WT and C543S caused significantly higher levels of cell migration in response to EGF, compared to C312S (Figure 2B). To determine if the increased cell migration was not limited to that specific cell line, an additional metastatic breast cancer cell line (triple negative breast cancer MDA-MB-468) was examined upon overexpressing ARHGEF7 WT or C312S in the presence or absence of EGF. Consistently, the same outcomes were observed (Figure S5C). MDA-MB-468 cells expressing ARHGEF7 WT exhibited significantly elevated cell migration in response to EGF, whereas cells expressing ARHGEF7 C312S had a slight increase in cell migration (Figure S5C).

**Figure 2.**
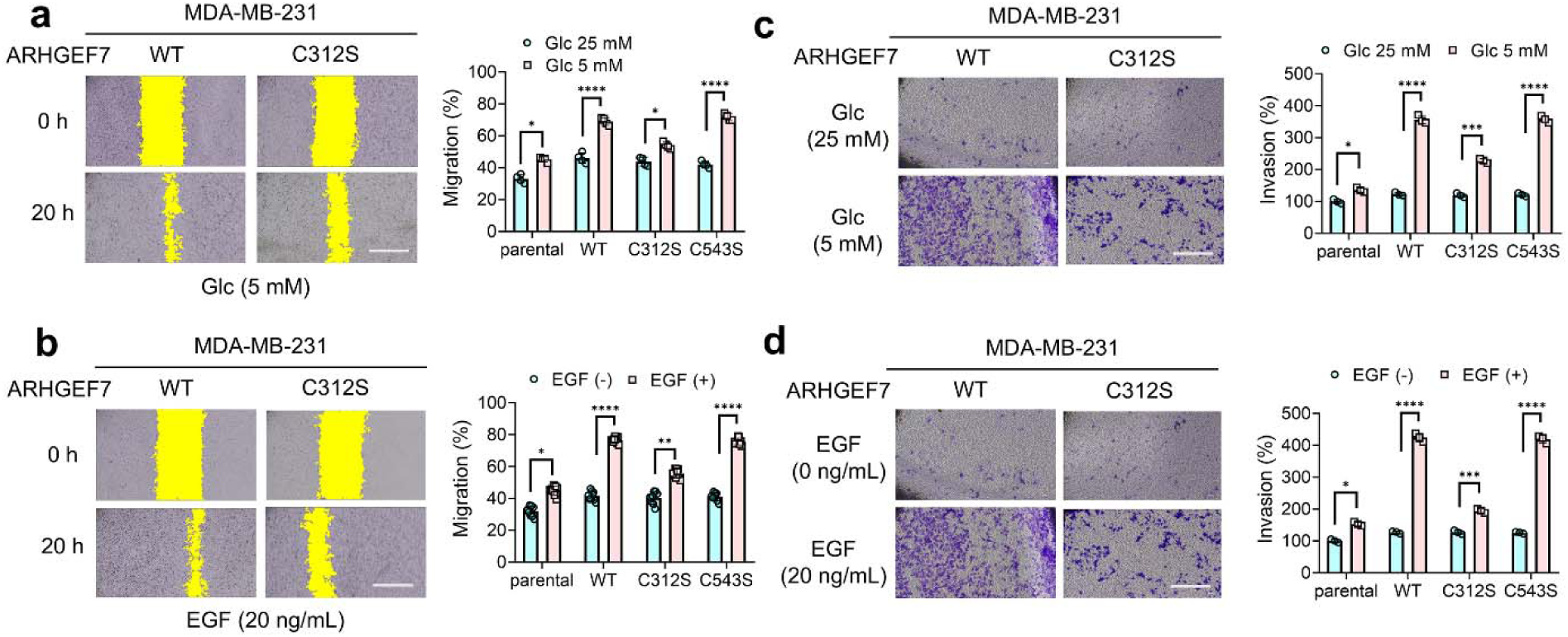
ARHGEF7 C312 glutathionylation increases cell migration and invasion. (**a**-**b**) *In-vitro* scratch migration analysis of MDA-MB-231 cells expressing ARHGEF7 construct in high glucose (HG, 25 mM) or low glucose (LG, 5 mM) conditions (n=4) (**a**) or in response to EGF (n=9) (**b**). Yellow indicates areas without cells. (**c**-**d**) Transwell invasion analysis of MDA-MB-231 cells expressing ARHGEF7 constructs in HL or LG (n=3) (**c**) or upon incubating EGF (n=3) (**d**). Cells were incubated in HG or LG (**a**, **c**) or with EGF for 20 h (**b**, **d**). A scale bar = 500 μm. Data represent the mean ± SD. The statistical difference was analyzed by two-way ANOVA and Tukey’s post-hoc test (**a**-**d**), where *p < 0.03, **p < 0.002, ***p < 0.0002, ****p < 0.0001.

In addition to migration, the effect of ARHGEF7 S-glutathionylation on cell invasion was investigated. MDA-MB-231 cells expressing ARHGEF7 WT, C312S, or C543S were grown in either HG or LG conditions and observed for cell invasion (Figure 2C). MDA-MB-231 cells expressing ARHGEF7 WT, along with C543S, showed dramatic increases in cell invasion in response to LG over HG (Figure 2C). In comparison, cells expressing C312S also caused increased invasion in LG (Figure 2C) but not as high as cells expressing WT or C543S. The same pattern was observed when MDA-MB-231 cells were stimulated with EGF. Cells expressing ARHGEF7 WT or C543S cells showed the largest increase in cell invasion, when compared to cells expressing C312S or non-transfected cells (Figure 2D). In agreement, MDA-MB-468 cells exhibited the same pattern as MDA-MB-231 cells when tested with EGF (Figure S5D). These data support that ARHGEF7 C312 S-glutathionylation increases cell migration and invasive phenotypes in response to EGF or oxidative stress.

### ARHGEF7 glutathionylation activates Rac1 and PAK1 pathways

The major biological function of ARHGEF7 is to bind and target Rac1 and PAK1 to cell protrusion or focal adhesions,^57,58^ where ARHGEF7 activates Rac1 via its GEF activity. The GTP-bound active Rac1 (Rac1-GTP) binds to PAK1, leading to PAK1 autophosphorylation and activation.^86^ Active PAK1 increases cell motility by phosphorylating and activating LIMK and its downstream targets, including cofilin, thereby increasing actin polymerization and dynamics.^61^ Active PAK1 also increases cell proliferation and migration through phosphorylating MEK1 to stimulate the MEK1/ERK1 pathway.^87^ The Rac1-PAK1 signaling cascade plays a major role in cell migration and is associated with cancer progression.^62^ Therefore, it was hypothesized that ARHGEF7 glutathionylation may modulate interactions with its protein partners, such as Rac1 and PAK1, or increase its enzymatic activity.

We monitored ARHGEF7 interactions with Rac1 and PAK1 in oxidative stress via co-immunoprecipitation (co-IP). MDA-MB-231 cells expressing ARHGEF7 WT or C312S were grown in HG or LG conditions. ARHGEF7 WT showed a significant increase in binding with Rac1 in LG versus HG (lane 2 vs. 1), whereas the increased binding was not observed in C312S in LG (lane 4 vs. 3, Figure 3A), suggesting that ARHGEF7 C312 glutathionylation increases its complex formation with Rac1. In contrast, both ARHGEF7 WT and C312S decreased binding to PAK1 (lanes 2 and 4 vs. 1 and 3, Figure 3A), supporting the idea that PAK1 dissociates from ARHGEF7 under oxidative stress (i.e., LG versus HG), but this dissociation is not attributed to ARHGEF7 C312 glutathionylation. Following the increased binding of ARHGEF7 with Rac1 upon C312 glutathionylation, we monitored the levels of active GTP-bound Rac1. MDA-MB-231 cells expressing ARHGEF7 WT showed a dramatic increase in GTP-bound active Rac1 in LG over HG (Figure 3B, lanes 2 vs. 1), which was completely abolished in cells expressing C312S (Figure 3B, lanes 4 vs. 3), suggesting that ARHGEF7’s increased binding to Rac1 upon its C312 glutathionylation leads to Rac1 activation. The observation is consistent with data showing that Rac1 associates with ARHGEF7 to promote its activation, whereas activated PAK1 dissociates from ARHGEF7 during cell migration.^57^

**Figure 3.**
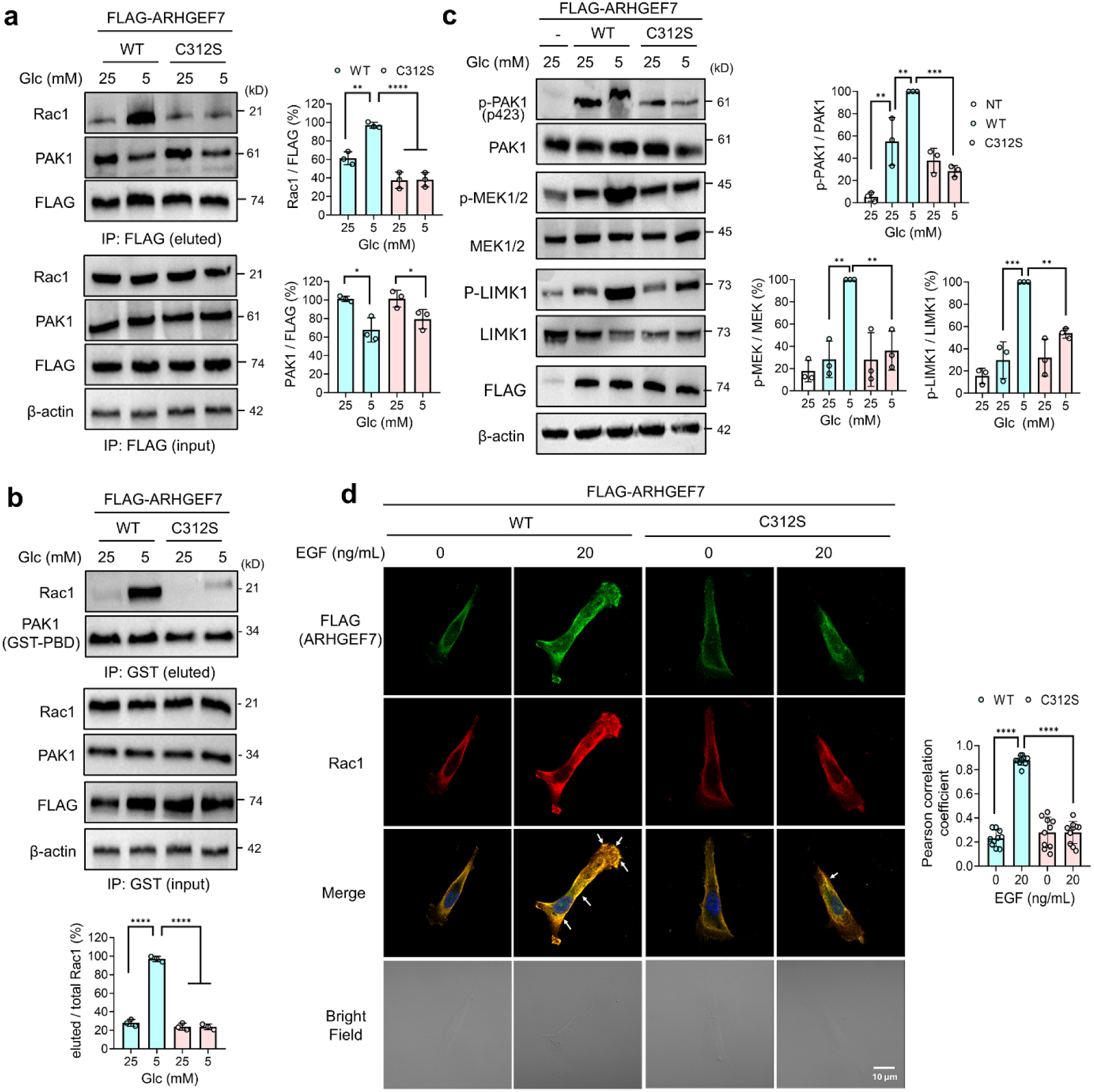
ARHGEF7 C312 glutathionylation activates the Rac1-PAK1 signaling cascade. Analysis of ARHGEF7 protein interactions and Rac1-PAK1 downstream signaling. MDA-MB-231 cells expressing ARHGEF7 WT or C312S were incubated in high glucose (HG, 25 mM) or low glucose (LG, 5 mM) for 20 h. (**a**) ARHGEF7 S-glutathionylation increases its binding to Rac1. ARHGEF7 co-immunoprecipitation (co-IP) with Rac1 and PAK1 (n=3). (**b**) Rac1 is activated upon ARHGEF7 S-glutathionylation. Active Rac1 was enriched by purified GST-PBD (p21-binding domain derived from PAK1) and analyzed by western blots (n=3). (**c**) ARHGEF7 S-glutathionylation activates PAK1 and its downstream signaling. Phosphorylation levels of PAK1, LIMK1, and MEK1/2 were analyzed by western blot (n=3). (**d**) ARHGEF7 S-glutathionylation increases Rac1 localization at the cell periphery. The co-localization images of ARHGEF7 and Rac1 (left). Cells were fixed and analyzed by antibodies to FLAG (green) and Rac1/Cdc42 (red) (n=10 images). White arrows indicate the intense co-localization of ARHGEF7 and Rac1. Pearson’s correlation coefficients (right) were calculated after Intermodes thresholding to select for colocalization of ARHGEF7 and Rac1 at the cell periphery (see also Figure S6). A scale bar = 10 μm. Data represent the mean ± SD. The statistical difference was analyzed by one-way ANOVA and Tukey’s post-hoc test (**a-d**), where *p < 0.03, **p < 0.002, ***p < 0.0002, ****p < 0.0001.

**Figure 4.**
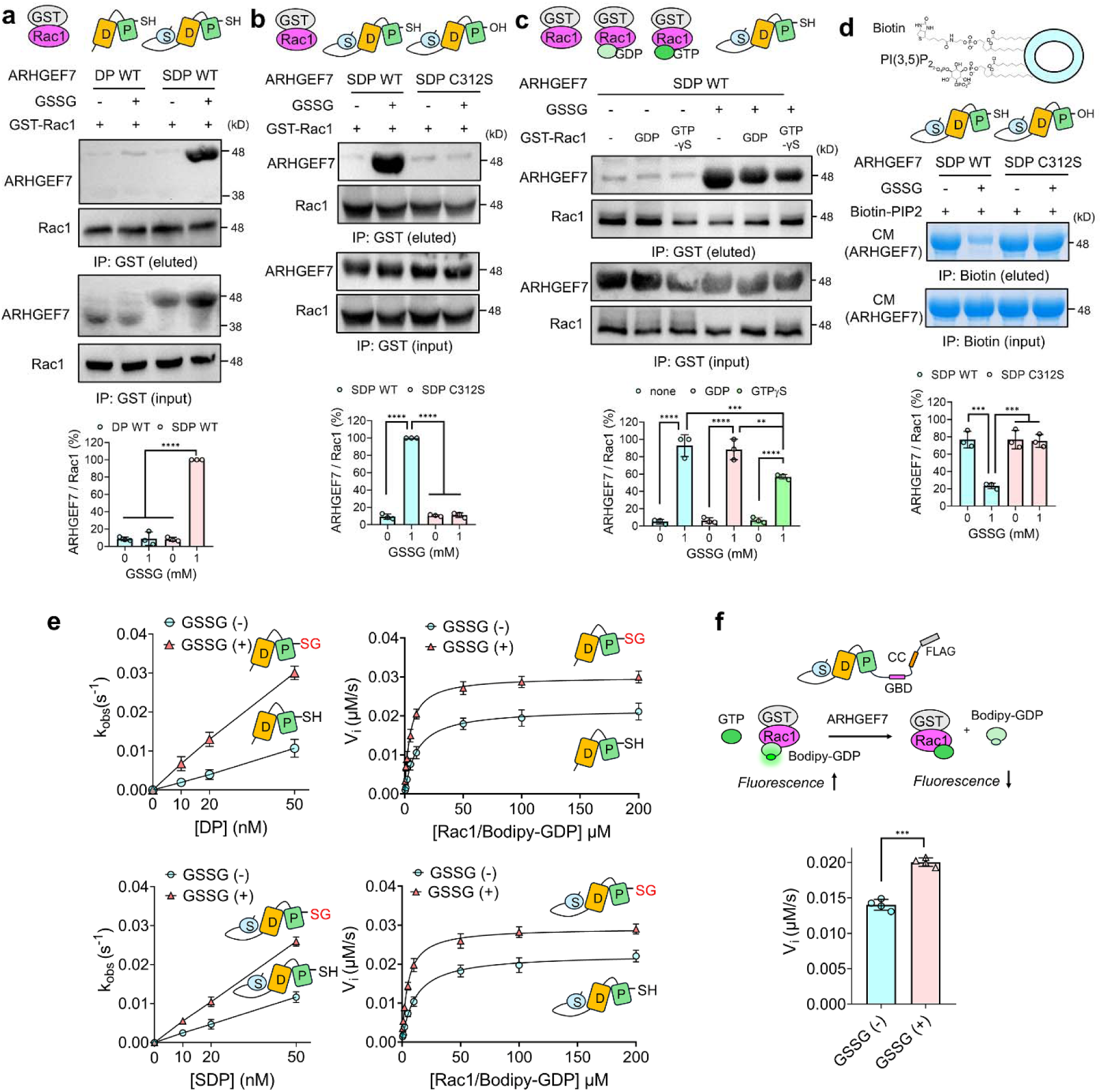
ARHGEF7 C312 glutathionylation increases its direct interactions with Rac1 and its GDP-GTP exchange activity for Rac1. (**a**-**c***) In vitro* binding analysis of ARHGEF7 constructs (DP or SDP) with GST-Rac1. Purified ARHGEF7 constructs (DP and SDP) were incubated with oxidized glutathione (GSSG) to induce S-glutathionylation. Following dialysis, ARHGEF7 constructs were incubated with GST-Rac1 immobilized on glutathione-Sepharose. GST-Rac1 and ARHGEF7 constructs were analyzed by Western blot before and after GST pull-down. (**a**) Comparison of Rac1 binding to ARHGEF7 DP WT and ARHGEF7 SDP WT (n=3). (**b**) Comparison of Rac1 binding to ARHGEF7 SDP WT and ARHGEF7 SDP C312S (n=3). (**c**). Comparison of ARHGEF7 SDP WT binding to Rac1 with different active site states (none, GDP, GTPγS) (n=3). (**d**) In vitro binding analysis of ARHGEF7 with PI(3,5)P_2_. Purified ARHGEF7 SDP treated without and with GSSG were incubated with biotinylated PI(3,5)P_2_ on micelle. Bound ARHGEF7 SDP constructs were analyzed by Coomassie stain (CM) (n=3). (**e**-**f**) ARHGEF7 increases its GDP-GTP exchange rate upon S-glutathionylation. Different purified ARHGEF7 constructs were incubated with GSSG and used for enzyme kinetics. Rates were determined by a loss of fluorescence when Bodipy-GDP bound to Rac1 dissociates from Rac1. (**e**) Rate measurement by purified ARHGEF7 DP or SDP constructs without or with incubation of GSSG. Rates were determined by increasing enzyme concentration (n=3) (left) and substrate concentration (Michaelis-Menten) (n=3) (right). (**f**) Rate measurement by a full-length ARHGEF7 construct without or with glutathionylation. FLAG-ARHGEF7 purified by affinity purification was incubated with GSSG and used to measure rates (n=4). Data represent the mean ± SD. The statistical difference was analyzed by one-way ANOVA and Tukey’s post-hoc test (**a-d**) or unpaired t-test (**f**), where *p < 0.03, **p < 0.002, ***p < 0.0002, ****p < 0.0001.

**Figure 5.**
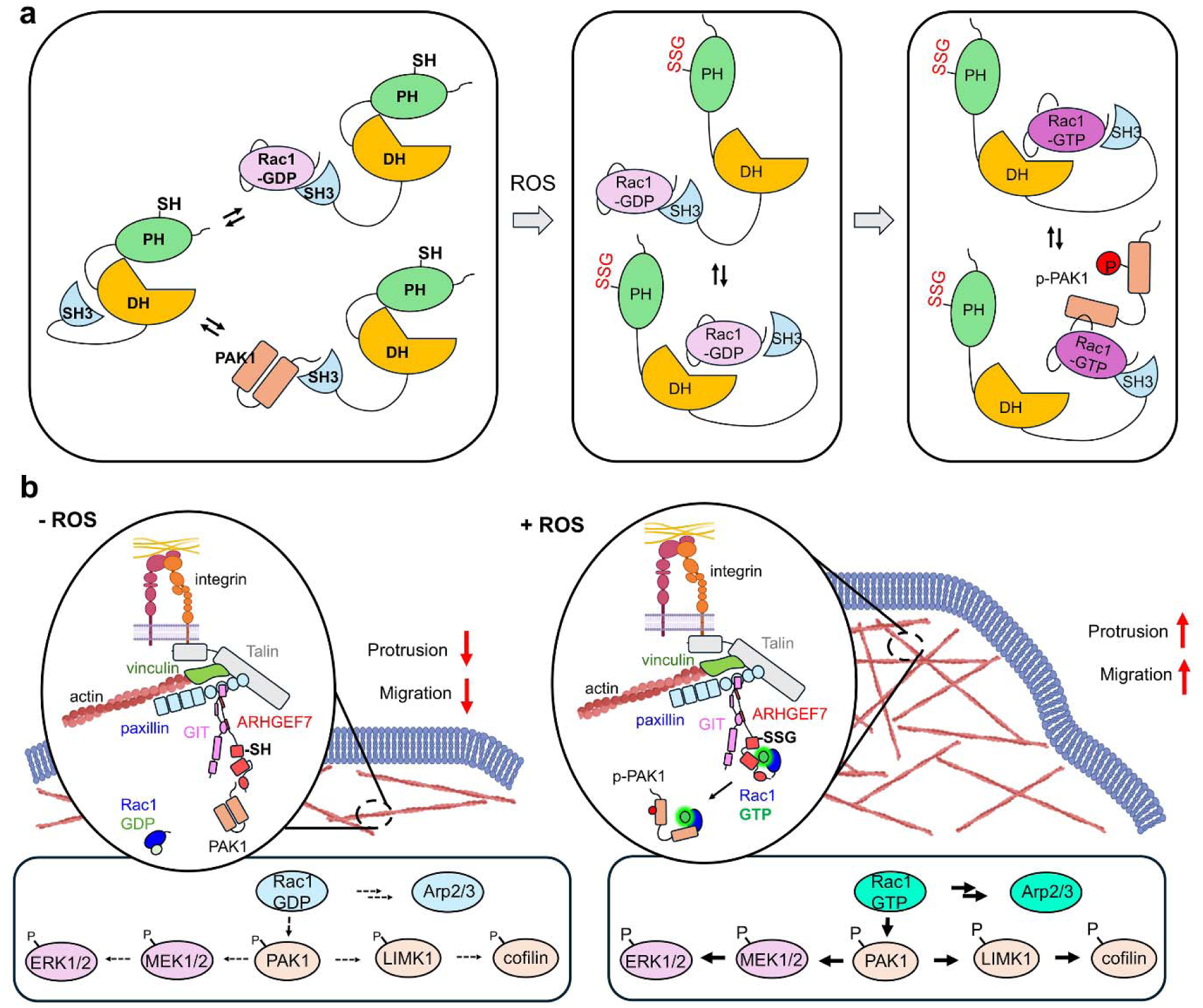
A model for increased cell migration by ARHGEF7 C312 glutathionylation. (**a**) A potential mode of ARHGEF7 activation by S-glutathionylation. In the absence of ROS, the DH-PH active site is relatively closed, and the SH3 domain stays bound to the DH-PH or its binding partners, Rac1 or PAK1, through the C-terminal proline-rich sequence (PRS) in Rac1 and PAK1 (left). Upon ROS generation, ARHGEF7 is glutathionylated at C312 at the PH domain, which opens the active site in the DH-PH domains to which SH3-bound Rac1-GDP binds for GDP-GTP exchange (middle). Rac1 activation follows, resulting in PAK1 phosphorylation and activation (right). (**b**) A model for ARHGEF7 activation in cells. In the absence of ROS, ARHGEF7, in complex with GIT1, may be associated with none or PAK1 in cells, without significant Rac1 activation. In the presence of ROS, glutathionylated ARHGEF7/GIT1/PAK1 complex is recruited to the nascent focal adhesion at the protrusion, where PAK1 is dissociated, and glutathionylated ARHGEF7 increases the binding to Rac1 for Rac1 activation. Active Rac1 increases phosphorylation and activation of PAK1. The active Rac1 and PAK1 activate their downstream signaling, including Arp2/3, LIMK-cofilin, and MEK1/2-ERK1, which increases branching and polymerization of actin filaments, lamellipodia formation, and cell migration.

Therefore, we examined the effects of ARHGEF7 glutathionylation on the Rac1-PAK1 downstream signaling pathways. Upon Rac1 activation, we examined its downstream cascade, including PAK1, LIMK1, and MEK1. The ectopic expression of ARHGEF7 WT or C312S constructs increased phosphorylation of PAK1 but less significantly on LIMK1 and MEK1/2, supporting ARHGEF7’s direct effects on PAK1 phosphorylation. However, MDA-MB-231 cells expressing ARHGEF7 WT showed increased phosphorylation and activation of PAK1, LIMK1, and MEK1/2 in LG versus HG (Figure 3C, lanes 3 vs 2), whereas ARHGEF7 C312S did not cause any significant changes in their phosphorylation (Figure 3C, lanes 5 vs. 4), suggesting the activation of the PAK1 downstream cascade upon ARHGEF7 C312 glutathionylation.

In addition to HG and LG conditions, the downstream effects of ARHGEF7 S-glutathionylation were monitored upon incubating with EGF, displaying the same outcome (Figure S5E-F). ARHGEF7 WT-expressing cells showed elevated binding with Rac1, compared to C312S, in response to EGF (Figure S5E). Subsequently, phosphorylation levels of PAK1 and its downstream effectors (i.e., LIMK1 and MEK1/2) were significantly elevated in WT-expressing cells upon incubating with EGF, compared to C312S-expressing cells (Figure S5F). These experiments demonstrate that ARHGEF7 S-glutathionylation activates Rac1 and its downstream signaling pathways, thereby promoting cell migration and invasion.

### ARHGEF7 glutathionylation increases recruitment of Rac1 to cell membrane and protrusions

In addition to activating Rac1 and PAK1, ARHGEF7 plays an important role in targeting Rac1 to focal adhesions and protrusions.^57,88^ At the protrusion, Rac1 promotes actin polymerization and stabilizes nascent focal adhesions to increase cell migration.^60^ Because ARHGEF7 glutathionylation increases the binding between ARHGEF7 and Rac1, we then monitored the recruitment and localizations of ARHGEF7 and Rac1 upon ARHGEF7 glutathionylation. MDA-MB-231 cells expressing ARHGEF7 WT or C312S were incubated without and with EGF, and cells were visualized by co-immunostaining analysis of ARHGEF7 (FLAG, green) and Rac1 (red). In the absence of EGF, ARHGEF7 WT and Rac1 were observed in the cytoplasm with diffused localizations (Pearson’s coefficient = 0.23 ± 0.07, n=10, Figure 3D, column 1, and Figure S6A). However, upon incubating EGF, cells increased apparent lamellipodia formation, where both intensity and localizations of ARHGEF7 WT and Rac1 were significantly enhanced in the membrane and protrusions (Pearson’s coefficient = 0.88 ± 0.04, n=10, Figure 3D, column 2, and Figure S6B). In contrast, cells expressing C312S did not change co-localizations or intensity of ARHGEF7 and Rac1 in the absence or presence of EGF (Pearson’s coefficient = 0.28 ± 0.1 and 0.28 ± 0.09, respectively, n=10, Figure 3D, column 3-4, and Figure S6C-D). These data support that ARHGEF7 C312 glutathionylation increases its binding to Rac1, thereby targeting Rac1 to the protrusion and facilitating lamellipodia formation.

### ARHGEF7 glutathionylation increases its direct interactions with Rac1

Our co-IP experiments support the idea that glutathionylated ARHGEF7 increases its binding to Rac1. To corroborate their direct enhanced association between glutathionylated ARHGEF7 and Rac1, recombinant ARHGEF7 and Rac1 constructs were analyzed for their binding *in vitro*. GST-Rac1 was used in a pull-down assay to determine Rac1 binding interactions with ARHGEF7 constructs [DH-PH (DP) and SH3-DH-PH (SDP)] pre-incubated with GSSG to induce glutathionylation. ARHGEF7 SDP, without glutathionylation, showed minimal binding to Rac1 (Figure 4A, lane 3). However, glutathionylated ARHGEF7 SDP showed strong binding interaction with Rac1 (Figure 4A, lanes 4 vs. 3). Interestingly, ARHGEF7 DP (i.e., DH-PH), without the SH3 domain, did not show any significant binding to Rac1 without or with glutathionylation (Figure 4A, lanes 2 and 1), suggesting that the SH3 is required for the enhanced binding interaction between glutathionylated ARHGEF7 and Rac1, which is consistent with the data that SH3 is responsible for binding to Rac1.^57^ To confirm the importance of C312 for glutathionylation, the same pull-down assay was performed to compare ARHGEF7 SDP WT and C312S. Unlike the strong binding of SDP WT with Rac1 upon SDP glutathionylation (Figure 4B, lanes 2 vs. 1), SDP C312S was unable to bind to Rac1 regardless of its glutathionylation state (Figure 4B, lanes 4 vs. 3). These results suggest that the SH3 domain may be inaccessible or sequestered by the DH-PH catalytic domains, being unable to bind Rac1 until disruption of the sequester by S-glutathionylation. The mechanism is analogous to other GEFs, such as Vav and ASEF, whose DH-PH domains are inhibited by their SH3 domains but activated by phosphorylation or protein-protein interactions.^89,90^

RhoGTPases, including Rac1, change their conformation, especially their effector loops, upon binding of GTP or GDP.^91,92^ Therefore, Rac1 active site occupancy was also examined for its effect on ARHGEF7 binding. Rac1 was incubated with none, GDP, or non-hydrolyzable GTPγS. The three Rac1 modes were examined for their binding to ARHGEF7 SDP WT. Regardless of Rac1 active site occupancy, ARHGEF7 SDP without glutathionylation minimally binds to Rac1 (Figure 4C, lanes 1-3). On the other hand, glutathionylated ARHGEF7 SDP showed high levels of binding to Rac1. Interestingly, non- or GDP-bound Rac1 showed higher binding to SDP than GTP-bound Rac1 (Figure 4C). Given that ARHGEF7 is used for GDP-GTP exchange in Rac1, the decreased binding of GTP-Rac1 versus GDP-Rac1 could facilitate its turnover via the product dissociation, as observed with other GEFs.^93^ These data further support the conclusion that ARHGEF7 C312 glutathionylation is responsible for the enhanced binding to Rac1, promoting its activation.

### ARHGEF7 glutathionylation disrupts PI(3,5)P_2_ binding *in vitro*

Phosphoinositides (PIPs) are important for focal adhesion dynamics and the regulation of cell migration, and they shuttle target proteins to lipid membranes.^94^ PIPs exhibit weak binding affinity for PH domains and have been proposed to induce conformational changes in DH-PH domains and enhance GEF activity.^95-97^ Previously, ARHGEF7 was shown to bind to PI(3,5)P_2_,^75^ with the predicted binding site in the PH domain at charged residues near C312, suggesting that PIP_2_ interaction with ARHGEF7 may be altered upon ARHGEF7 glutathionylation. Therefore, a pull-down assay of ARHGEF7 SDP with or without glutathionylation was performed using biotinylated PI(3,5)P_2_ beads. ARHGEF7 SDP WT dramatically decreased its binding to PI(3,5)P_2_ upon SDP glutathionylation (Figure 4D, lanes 2 vs. 1). On the other hand, SDP C312S did not change its binding interaction with PI(3,5)P_2_ (Figure 4D, lanes 4 vs. 3), supporting that ARHGEF7 C312 glutathionylation interferes with PIP_2_ binding. To our knowledge, the functional significance of ARHGEF7 and PI(3,5)P_2_ interactions has not been demonstrated. However, PI(3,5)P_2_ is known to transiently increase its level, especially at the endosome, under stress.^76,98^ Therefore, it is possible that PI(3,5)P_2_ may regulate the localization of ARHGEF7 at the early time points under stress, which could be displaced after ARHGEF7 glutathionylation.

### ARHGEF7 glutathionylation increases the Rac1 GDP-GTP exchange activity

PIP_2_ was shown to bind to the DH-PH domain in the RhoGEF and increase its GEF activity.^97^ Because the glutathionylation site is located at the PIP_2_ binding site, we next evaluated any change in ARHGEF7 GEF enzymatic activity upon C312 glutathionylation. To measure GEF activity, ARHGEF7 DP and SDP were glutathionylated by GSSG, while GST-Rac1 was incubated with fluorescent BODIPY-GDP (Rac1/BODIPY-GDP as a substrate for GEF). ARHGEF7 enzyme activity was monitored via a loss of fluorescence that occurs upon displacement of fluorescent GDP in the presence of excess GTP (Figure 4F).^99^

To begin with, initial velocities were measured by using a saturating amount of Rac1/BODIPY-GDP and a series of ARHGEF7 enzyme concentrations. Without glutathionylation, ARHGEF7 DP (k_cat_/K_M_ = 2.2 x 10^5^ M^-1^s^-1^) and ARHGEF7 SDP (k_cat_/K_M_ = 2.3 x 10^5^ M^-1^s^-1^) showed very similar exchange rates (Figure 4E, left). However, after incubation with GSSG, both ARHGEF7 DP (k_cat_/K_M_ = 6.0 x 10^5^ M^-1^s^-1^) and SDP (k_cat_/K_M_ = 5.2 x 10^5^ M^-1^s^-1^) increased their exchange rates (Figure 4E, left). Notably, both DP and SDP showed similar rates before and after glutathionylation, with a 2-3-fold increase in the exchange rate upon glutathionylation (Figure 4E).

To corroborate the data, Michaelis-Menten kinetics were performed to measure catalytic efficiency using varying concentrations of Rac1/BODIPY-GDP. Consistently, the identical data were obtained, showing the similar catalytic efficiencies of ARHGEF7 DP (V_max_ = 0.022 s^-1^, K_M_ = 10.19 μM, k_cat_/K_M_ = 2.14 x 10^5^ M^-1^s^-1^) and SDP (V_max_ = 0.023 s^-1^, K_M_ = 11.01 μM, k_cat_/K_M_ = 2.05 x 10^5^ M^-1^s^-1^) (Figure 4E, right). Importantly, glutathionylation of ARHGEF7 increased its catalytic efficiencies, when compared to non-glutathionylated constructs, but to the comparable values between ARHGEF7 DP (V_max_ = 0.030 s^-1^, K_M_ = 4.57 μM, k_cat_/K_M_ = 6.58 x 10^5^ M^-1^s^-1^) and SDP (V_max_ = 0.029 s^-1^, K_M_ = 4.81 μM, k_cat_/K_M_ = 6.08 x 10^5^ M^-1^s^-1^) (Figure 4E, right). Glutathionylation of both ARHGEF7 constructs activated their GEF activity by increasing V_max_ and decreasing K_M_ values, suggesting that glutathionylation of the PH domain likely induces conformational change in DH-PH, enabling the DH domain to increase Rac1 binding and turnover.

Finally, full-length ARHGEF7 was assayed to validate the observed enhancement in the GEF activity upon its glutathionylation. MDA-MB-231 cells expressing ARHGEF7 WT were collected. Full-length ARHGEF7 protein was purified through FLAG-affinity purification (Figure S7). Full-length ARHGEF7 was incubated with and without GSSG to induce glutathionylation. Full-length ARHGEF7 showed a significant increase in the GDP-GTP exchange rate with Rac1 after incubating GSSG (Figure 4F). These data support that ARHGEF7 C312 glutathionylation increases the DH-PH’s GEF activity for Rac1.

These data support our model that at the molecular level (Figure 5A), in the absence of ROS, the SH3-DH-PH domain of ARHGEF7 may have a conformation where the DH-PH active site is relatively closed and the SH3 is sequestered to the DH-PH or capable of binding to PAK1 and inactive Rac1-GDP in competition, which harbor the C-terminal proline residues responsible for binding to the SH3 domain. Upon ROS generation, ARHGEF7 is glutathionylated at C312 in the PH domain. Glutathionylation in the PH domain opens the active site in the DH-PH domains, enabling the SH3-bound Rac1-GDP to access the DH domain for GDP-GTP exchange and Rac1 activation. At the cell level (Figure 5B), in the absence of ROS, ARHGEF7, in complex with GIT1, may associate with PAK1 at focal adhesions in cells without significant Rac1 activation. However, in the presence of ROS, the ARHGEF7/GIT1/PAK1 complex is recruited to the nascent focal adhesion at the protrusion, where PAK1 dissociates. Glutathionylated ARHGEF7 increases its binding to Rac1 for its activation, along with PAK1 activation and phosphorylation. The active Rac1 and PAK1 then increase downstream signaling, including Arp2/3, LIMK1-Cofilin, and MEK1/2-ERK1, thereby promoting actin polymerization, lamellipodia formation, and cell migration.

## Discussion

Numerous studies have demonstrated that ROS play a key role in regulating biological processes of cell migration, adhesion, invasion, epithelial-mesenchymal transition (EMT), and cancer metastasis.^6,7^ Multiple signaling pathways have been modulated in response to ROS, including activation of mitogen-activated protein kinase (MAPK)^10^ and other kinases (e.g., Src, MEKK1, FAK, and PKC),^6,48,100^ inhibition of protein tyrosine phosphatases (e.g., PTP),^101^ dynamics of cytoskeletal filament formation (e.g., actin, cofilin),^102^ dissociation or modulation of cell-cell adhesion (e.g., integrin, paxillin, and p120),^34,103^ and degradation of extracellular matrix proteins (e.g., increased MMP).^7^ Among them, redox regulation of the Ras superfamily and of RhoGTPases (Rac1 and RhoA), which are oncogenic signaling hubs, has been implicated in cell proliferation, migration, and adhesion.^104^ Nevertheless, functional analyses of redox-sensitive cysteines and their direct roles in cell migration, invasion, and cancer metastasis have been limitedly demonstrated.

In this report, we showed that RhoGEF ARHGEF7 is susceptible to S-glutathionylation, which increases cell migration and invasion by activating the Rac1 GTPase, a central player in cell migration and invasion. Despite the presence of 10 cysteines in ARHGEF7, we found that glutathionylation occurs prominently at C312 in response to EGF, which induces ROS production and promotes cell migration. Similarly, ARHGEF7 glutathionylation is selectively induced at C312 by limited glucose levels that cause oxidative stress, a condition that has also been shown to increase cell migration and invasion.^31,32^ C312 is relatively exposed to the surface, surrounded by basic residues in the PH domain. Although many other cysteines in ARHGEF7 are predicted to have higher surface exposures (i.e., ASA) and lower pK_a_ values, it is plausible that the PH domain may have a glutathione-binding or interacting site that mediates S-glutathionylation, thus contributing to the selectivity at C312. Additional structural analyses may be necessary to further support the presence of a potential glutathione-binding site or glutathionylation selectivity in the PH domain.

The biochemical effects of S-glutathionylation in ARHGEF7 are two-fold. First, S-glutathionylation increases the ARHGEF7 GEF activity. Although the exact mechanism of GEF activation remains to be elucidated, the mode of activation can be inferred from the cysteine residue, C312, targeted by S-glutathionylation. C312 is located in the PH domain of ARHGEF7. PH domains are invariably found in tandem DH-PH domains in the RhoGEF family,^56^ where PH inhibits or regulates the DH’s GEF activity, participates in binding to GTPases, or acts as a membrane anchor via binding to phospholipids.^71^ For example, phospholipid binding to the PH domain was shown to enhance GEF activity by disrupting interactions between the PH domain and the catalytic DH domain.^95-97^ C312 is positioned at the predicted phospholipid binding site, and ARHGEF7 C312 glutathionylation interferes with PI(3,5)P_2_ binding while increasing ARHGEF7 GDP-GTP exchange activity. Based on this data, it is plausible to hypothesize that upon ARHGEF7 glutathionylation, glutathione occupies the phospholipid-binding sites in the PH domain, which induces a DH-PH conformational change, as phospholipids do,^97^ thereby increasing the binding and turnover of a Rac1 substrate. Second, ARHGEF7 C312 glutathionylation significantly increases its binding to Rac1. It was reported that both Rac1 and PAK1 contain C-terminal proline-rich sequences (PRS) that are responsible for their competitive binding to the SH3 domain of ARHGEF7.^57^ In our data, Rac1 binding is significantly higher with glutathionylated SDP than with glutathionylated DP, suggesting that the SH3 domain is required for Rac1 binding. However, non-glutathionylated SDP did not display significant binding to Rac1, compared to glutathionylated SDP, suggesting that the SH3 domain may be inaccessible to the Rac1 PRS, potentially by binding to the DH-PH domain.^90^ However, the SH3 is unlikely to block the active site of ARHGEF7 in the DH domain because no major differences in GEF activity were observed in the Michaelis-Menten kinetics between SDP and DP (or their glutathionylated couterparts). Regardless, glutathionylation may release the SH3 and DH-PH interactions, enabling SH3 to bind Rac1. Alternatively, the SH3 domain in non-glutathionylated SDP may be available to bind Rac1, but not as highly as after glutathionylation. Interestingly, our data also showed that Rac1-GDP binds to glutathionylated SDP higher than Rac1-GTP. Because Rac1-GDP and Rac1-GTP have different conformations, especially in a switch II loop that interacts with the active sites in GEFs,^91,92^ such as the DH-PH domains in ARHGEF7, the differential binding affinity between Rac1-GDP and Rac1-GTP suggests that the DH-PH domains in glutathionylated ARHGEF7 likely contribute to the enhanced binding to Rac1. Therefore, the observed high Rac1 binding to ARHGEF7 is likely due to both glutathionylated DH-PH and SH3 domains interacting synergistically or additively with Rac1. Eventually, the SH3 in glutathionylated ARHGEF7 could keep Rac1-GDP in close proximity and recycle it to active Rac1-GTP, thereby maintaining Rac1-GTP in an active state for a longer period. Overall, ARHGEF7 S-glutathionylation would sustain active Rac1 signaling by increasing its binding to Rac1 and its GEF enzymatic activity.

Upon ARHGEF7 S-glutathionylation, the activated Rac1 was shown to activate PAK1 and its downstream signaling, including LIMK1-cofilin and MEK1-Erk1/2 axes, which promote actin polymerization at the protrusion.^61,62^ Because Rac1/PAK1 activation is sufficient to induce signaling cascades for cell migration, invasion, and metastasis,^62,105^ our report uncovers a potential redox mechanism (i.e., ARHGEF7 S-glutathionylation) that contributes to metastatic cancer progression, consistent with higher ARHGEF7 expression in many cancers,^64-67^ including breast cancer, which remains to be validated in the future. In addition to cancer cells, ARHGEF7 is highly expressed in neuronal cells, where it activates Rac1/PAK1, promoting axon formation and neuronal migration.^106-108^ Overexpression of ARHGEF7 has been shown to improve dendritic and synaptic integrity, potentially reversing cognitive deficits.^109^ Therefore, it will be interesting to see the effect of ARHGEF7 S-glutathionylation in other cell types.

In this report, we mainly focused on SH3-DH-PH domains due to their proximity to C312, the primary site of S-glutathionylation. In addition to C312, our proteomic analysis identified C543, located in the α-helical GBD domain and relatively proximal to the binding interface between the GBD and the GIT protein (Figure S2C). However, our data shows that glutathionylation primarily occurs at C312, suggesting that glutathionylation at 543 may be less significant. ARHGEF7 and GIT1 form a trimer and a dimer, respectively, via oligomerization of their CC domains, while the ARHGEF7 GBD interacts with the GIT SHD domains, thereby forming a multimeric complex.^79^ Because of the relatively distant location of C312 away from the GBD and CC domains, ARHGEF7 and GIT complex formation is unlikely to be affected by ARHGEF7 C312 glutathionylation. However, GIT is important for the formation of focal adhesions with ARHGEF7.^77^ Further analysis will help clarify the implications of GIT in cell migration induced by ARHGEF7 C312 glutathionylation.

## Materials and Methods

### Molecular Cloning

FLAG-tagged mouse ARHGEF7 expression plasmid was obtained from Addgene (FLAG-betaPIXa, pcDNA3-FLAG-mARHGEF7 Cat# 15234). Cysteine to serine substitution mutants of mouse ARHGEF7 were generated through site-directed mutagenesis using the following forward primers and their reverse complements: C312S (5’-C CTG ATT CAG <u>TCT</u> GCC GGA AGT G -3’) and C543S (5’-C ATT GAA GCT TAC <u>AGC</u> ACC AGC GCC - 3’). Human cDNA clones of PAK1 binding domain (GST-PBD, pGEXTK-PAK1 70-117, Cat# 12217) and Rac1 (GST-Rac1, pGEX-Rac1, Cat# 12200) were obtained from Addgene. Additionally, specific domains of the mouse ARHGEF7 cDNA clone (NM_017402, Addgene, Cat# 15234, pcDNA3-FLAG-mARHGEF7) were subcloned into a pET28(+) bacterial expression vector using the NdeI and EcoRI restriction enzyme sites. The PCR product containing the DH and PH domains (ARHGEF7 DP) was generated with the forward primer (5’-GCT AGC <u>CAT ATG</u> TAT TAC AAC GTG G -3’, NdeI) and the reverse primer (5’-GAG CTC <u>GAA TTC</u> <u>TTA</u> CTT CGT CTG CTT C -3’, EcoRI, stop codon). The PCR product containing the SH3, DH, and PH domains (ARHGEF7 SDP) was generated with the forward primer (5’-CTA GCA CAT ATG ACT GAT AAC ACC AAC AGC CAA CTG -3’, NdeI) and the same reverse primer used to make ARHGEF7 DP. The PCR products and empty pET28(+) vector were double-digested with NdeI and EcoRI restriction enzymes, purified from an agarose gel, and ligated using T4 DNA ligase to create ARHGEF7 DP and ARHGEF7 SDP. The PCR product containing only the SH3 domain (ARHGEF7 SH3) was generated through site-directed mutagenesis of a tyrosine to a stop codon in between the SH3 and DH domains of ARHGEF7 SDP using the forward primer (5’-C AAC AAG AGC <u>TAG</u> TAC AAC GTG GTG C -3’, stop codon) and its reverse complement. All cloned plasmids were confirmed by DNA sequencing.

### Purification of Proteins

All proteins were transformed, expressed, and purified in BL21-CodonPlus(DE3)-RIL competent cells. Cells were grown in LB medium (1 L) at 37°C until OD_600_ was reached. Protein expression was induced through IPTG (1 mM) and incubated at 18°C for 20 h. Cells were harvested through centrifugation at 4,000 rpm for 15 min at 4°C. 6x-His-tagged proteins (ARHGEF7 DP, ARHGEF7 SDP, ARHGEF7 SH3) were resuspended in a lysis buffer (25 mM Tris, pH 7.4, 100 mM NaCl, 1 mM PMSF) and lysed by double passage through a chilled French press at 1,500 psi. Cell debris was pelleted by centrifugation at 13,000 rpm for 30 min at 4°C. The supernatant was incubated with HisPur™ Ni-NTA resin (Thermo Scientific) for 1 h at 4°C. The resin was washed three times with a low imidazole wash buffer (25 mM Tris, pH 7.4, 100 mM NaCl, 25 mM imidazole). The proteins were eluted in a high-imidazole elution buffer (25 mM Tris, pH 7.4, 100 mM NaCl, 250 mM imidazole), and the purity of the eluate was visualized on an SDS-PAGE gel. Pure fractions were collected and dialyzed overnight in a storage buffer (25 mM Tris, pH 7.4, 100 mM NaCl, 1 mM DTT).

GST-tagged proteins (GST-PBD, GST-Rac1) were resuspended in a lysis buffer (50 mM Tris, pH 8.0, 150 mM NaCl, 0.1 mM EDTA) and lysed by double passage through a chilled French press at 1,500 psi. Cell debris was pelleted by centrifugation at 13,000 rpm for 30 min at 4°C. The supernatant was incubated with Glutathione Sepharose™ 4 Fast Flow resin (Cytiva) for 1 h at 4°C. The resin was washed three times with the lysis buffer. The proteins were eluted in a glutathione elution buffer (50 mM Tris, pH 8.0, 150 mM NaCl, 0.1 mM EDTA, 10 mM reduced glutathione), and the eluates were analyzed on an SDS-PAGE gel. Pure fractions were collected and dialyzed overnight in a storage buffer (50 mM Tris, pH 8.0, 150 mM NaCl, 1 mM DTT).

### Cell Culture

Breast cancer cell line MDA-MB-231 (ATCC, HTB-26™) and breast cancer cell line MDA-MB-468 (ATCC, HTB-132™) were maintained in Dulbecco’s Modified Eagle Medium with high glucose (DMEM, Cytiva, Hyclone™) supplemented with 10% Fetal Bovine Serum (FBS, Cytiva, Hyclone™) and 1% (100 U/mL) Penicillin-Streptomycin (Pen-strep, Thermo Scientific) in a humified incubator at 37°C in a 5% CO_2_ environment.

### Transfection

Cells were transfected using Lipofectimine™ 3000 Reagent (Thermo Scientific) at 80% confluency following the manufacturer’s protocol. 10 µg ARHGEF7 plasmids (WT, C312S, or C543S), 15 µL P3000™ Reagent, and 20 µL Lipofectamine 3000™ Reagent were mixed (1 µg plasmid: 2 µL P3000: 1.5 µL Lipofectamine) before being added to each 10 cm dish of cells in Opti-MEM™ medium (Thermo Scientific). After incubating for 6 h, the Opti-MEM™ medium was replaced with DMEM supplemented with 10% FBS and 1% Pen-strep.

### Glutathionylation Analysis in Cells

MDA-MB-231 cells were transduced with an adenovirus (Ad-GS M4, Vector Biolabs) containing a modified glutathione synthetase (GS M4, F152A/S151G).^81^ 6 x 10^6^ PFU Ad-GS M4 and 5 µg Polybrene (Sigma Aldrich) were mixed and then added to each 10 cm dish of cells in DMEM supplemented with 2% FBS. After 6 h, the medium was replaced with DMEM supplemented with 10% FBS and 1% Pen-strep. Cells were subsequently transfected with pcDNA3-FLAG-ARHGEF7 (WT, C312S, or C543S). The following day, cells were supplemented with 0.6 mM azido-Ala to produce azido-glutathione. After 24 h, glutathionylation was induced in the cells by lowering glucose levels through changing the medium to glucose-free DMEM supplemented with different glucose concentrations (and 0.3 mM azido-Ala). Alternatively, glutathionylation was induced by adding epidermal growth factor (EGF) through changing the medium to DMEM supplemented with 20 ng/mL EGF (and 0.3 mM azido-Ala). After 20 h, the cells were collected in RIPA lysis buffer [50 mM Tris, pH 8.0, 150 mM NaCl, 1 mM EDTA, 1% NP-40, 0.25% sodium deoxycholate, 0.1% sodium dodecyl sulfate (SDS), protease inhibitor cocktail] containing 50 mM N-ethylmaleimide (NEM). Ice-cold acetone (4x volume) was used to precipitate 1 mg of cell lysate at -20°C for 30 min. After centrifugation at 8,000 rpm for 5 min, the pellet was resuspended in click reaction buffer (1x PBS, pH 7.4, 0.2% SDS) through sonication. Cu(I)Br dissolved in DMSO/tBuOH (3:1, v/v) was mixed with THPTA and then added to the resuspended lysate, followed by addition of biotin-alkyne (Vector Biolabs). The click reaction (1x PBS, pH 7.4, 0.2% SDS, 2 mM Cu(I)Br, 2 mM THPTA, and 400 µM biotin-alkyne) was incubated at room temperature for 1 h. The proteins were then precipitated in ice-cold acetone at -20°C for 30 min. After centrifugation at 8,000 rpm for 5 min, the pellet was resuspended in click reaction buffer. The resuspended proteins were added to a solution containing high-capacity streptavidin agarose beads (Pierce™) and rotated at 4°C overnight. The beads were pelleted through centrifugation at 1,400 rpm for 3 min and then washed three times with click reaction buffer. To elute the proteins, the beads were resuspended in 50 µL SDS-loading dye (2x) containing 10 mM DTT and incubated at 95°C for 10 min. Following centrifugation at 13,000 rpm for 3 min, the eluted proteins were separated by SDS-PAGE and transferred to the PVDF membrane for western blot analysis. The membrane was blocked in 5% BSA in TBST (1x TBS, pH 7.4, 0.1% Tween® 20) at room temperature for 30 min. The membrane was incubated with primary antibody to FLAG (1:1000, Sigma, Cat# F1804) or actin (1:1000) in 1% BSA in TBST at 4°C overnight. The membrane was washed three times in TBST for 10 min and then incubated with the corresponding horseradish peroxidase (HRP)-conjugated secondary antibodies in TBST at room temperature for 1 h. After washing three times in TBST for 10 min, the proteins on the membrane were visualized by enhanced chemiluminescence (SuperSignal™ West Pico PLUS) using the iBright imaging system (Thermo Fisher Scientific).

### Glutathionylation analysis of purified proteins in vitro

100 µg of purified ARHGEF7 DP or ARHGEF7 SDP was incubated with 1 mM azido-glutathione and a gradient of H_2_O_2_ (0-200 µM) at room temperature for 15 min. The reaction was incubated with 20 mM NEM at room temperature for 20 min to block remaining cysteines. Ice-cold acetone (4x volume) was used to precipitate the reaction at -20°C for 30 min. After centrifugation at 8,000 rpm for 5 min, the pellet was resuspended in click reaction buffer (1x PBS, pH 7.4, 0.2% SDS) through sonication. The click reaction (1x PBS, pH 7.4, 0.2% SDS, 2 mM Cu(I)Br, 2 mM THPTA, 100 µM TAMRA-alkyne) was incubated at room temperature for 1 h. The proteins were precipitated in ice-cold acetone at -20°C for 30 min. After centrifugation at 8,000 rpm for 5 min, the pellet was resuspended in click reaction buffer. After running on an SDS-PAGE gel, proteins were analyzed by Coomassie stain and fluorescence using the iBright imaging system (Thermo Scientific).

### Migration Assay

MDA-MB-231 and MDA-MB-468 cells (1 x 10^5^ cells) were seeded into a 12-well plate and grown to 100% confluency. After 3 h of serum starvation, a wound was created in the wells using a 10 µL micropipette tip. Detached cells were removed through three washes of the wells with warm PBS. Cells were incubated in glucose-free DMEM supplemented with either 25 mM or 5 mM glucose for 20 h. Alternatively, cells were incubated in DMEM supplemented with or without 20 ng/mL EGF. The wound was imaged at the start (0 h) and end (24 h) time points, with wound closure analyzed using an MRI wound-healing plugin in ImageJ.

### Invasion Assay

MDA-MB-231 and MDA-MB-468 cells (5 x 10^5^ cells) were seeded on top of a transwell insert coated with Matrigel (Fisher Scientific) in 12-well plate. Cells were incubated in glucose-free DMEM supplemented with either 25 mM or 5 mM glucose for 20 h. The lower chamber was supplemented with 10% FBS to act as a chemoattractant. Alternatively, cells were incubated in DMEM for 20 h with the lower chamber supplemented with 20 ng/mL EGF as a chemoattractant. The cells were washed three times with PBS followed by using a cotton swab on the top of the membrane to remove non-invasive cells. The cells on the bottom of the membrane were fixed in 4% formaldehyde at room temperature for 15 min and then permeabilized with PBST (PBS, pH 7.4, 0.1% Triton X-100) at room temperature for 15 min. After a wash with PBS, the cells were stained with a 0.2% (w/v) crystal violet solution at room temperature for 15 min. Excess dye was removed through three washes with PBS before the membranes were imaged on a microscope. A 33% acetic acid solution was then used to extract crystal violet dye, and the absorbance of the dye was measured at 560 nm using a Synergy H1 microplate reader (BioTek).

### Western blot

To analyze phosphorylation, MDA-MB-231 cells were transfected with ARHGEF7 WT or C312S and grown for 24 h. Cells were then incubated in glucose-free DMEM supplemented with 25 mM or 5 mM glucose for 24 h or 20 ng/mL EGF for 20 h. The cells were lysed in a Co-IP lysis buffer (50 mM Tris, pH 8.0, 150 mM NaCl, 1 mM EDTA, 1% NP-40, 0.25% sodium deoxycholate, protease inhibitor cocktail) with a PhosSTOP™ phosphatase inhibitor tablet (Roche). Proteins were separated by SDS-PAGE and transferred to the PVDF membrane. The membrane was blocked in 5% BSA in TBST at room temperature for 30 min. The membrane was incubated with primary antibody solutions, FLAG (1:1000, Sigma, Cat# F1804), phospho-PAK1 (1:1000, Cell Signaling, Cat# 2601S), PAK1 (1:1000, Cell Signaling, Cat# 2608S), phospho-LIMK1 (1:1000, Cell Signaling, Cat# 3841S), LIMK1 (1:1000, Cell Signaling, Cat# 3842S), phospho-MEK1 (1:1000, Cell Signaling, Cat# 9154S), MEK1/2 (1:1000, Cell Signaling, Cat# 9122S), actin (1:1000), in 1% BSA in TBST at 4°C overnight. The membrane was washed three times in TBST for 10 min and then incubated with the corresponding horseradish peroxidase (HRP)-conjugated secondary antibodies, anti-mouse (1:3000, Cytiva, NA931) or anti-rabbit (1:3000, Cytiva, NA934) in TBST at room temperature for 1 h. After washing three more times in TBST for 10 min, the proteins on the membrane were visualized by enhanced chemiluminescence (SuperSignal™ West Pico PLUS) using the iBright imaging system (Thermo Scientific).

### Co-Immunoprecipitation

MDA-MB-231 cells were transfected with FLAG-ARHGEF7 WT or C312S. After 24 h, cells were incubated in glucose-free DMEM supplemented with either 25 mM or 5 mM glucose for 24 h or 20 ng/mL EGF for 20 h. The cells were then lysed in a Co-IP lysis buffer. 1 mg of cell lysate was incubated with FLAG primary antibody (1:500) at 4°C for 1 h. The solution was added to a slurry of protein G agarose beads (Pierce™) and rotated at 4°C overnight. The beads were pelleted through centrifugation at 1,400 rpm for 3 min and then washed three times with PBS. To elute the proteins, the beads were resuspended in 50 µL SDS-loading dye (2x) containing 10 mM DTT and incubated at 95°C for 10 min. Following centrifugation at 13,000 rpm for 3 min, the eluted proteins were separated by SDS-PAGE and transferred to the PVDF membrane for western blot analysis, using FLAG (1:1000), PAK1 (1:1000), Rac1 (1:1000, Cell Signaling, Cat# 4651S), or actin (1:1000) in 1% BSA in TBST at 4°C overnight. The corresponding horseradish peroxidase (HRP)-conjugated secondary antibodies were used to visualize proteins by chemiluminescence. Blots were analyzed using the iBright imaging system (Thermo Scientific).

### Active Rac1 analysis

MDA-MB-231 cells were transfected with FLAG-ARHGEF7 (WT or C312S) and grown for 24 h. Cells were incubated in glucose-free DMEM supplemented with either 25 mM or 5 mM glucose for 24 h. The cells were then lysed in a Co-IP lysis buffer. 1 mg of cell lysate was incubated with 100 µg GST-PBD at 4°C for 1 h. The solution was added to Glutathione Sepharose™ 4 Fast Flow resin (Cytiva) and rotated at 4°C overnight. The beads were pelleted through centrifugation at 1,400 rpm for 3 min and then washed three times with PBS. Proteins were eluted in 50 µL SDS-loading dye (2x) containing 10 mM DTT. Proteins were separated by SDS-PAGE and transferred to the PVDF membrane for western blot analysis, using FLAG (1:1000), PAK1 (1:1000), Rac1 (1:1000), or actin (1:1000).

### Immunostaining

MDA-MB-231 cells transfected with FLAG-ARHGEF7 WT or C312S (1 x 10^4^ cells) were seeded into 35-cm plates with a coverslip (Mattek) that had been coated with Fibronectin. 48 h post-transfection, cells were incubated in DMEM supplemented with or without 20 ng/mL EGF for 24 h. After washing with PBS, the cells were fixed with 4% paraformaldehyde, pH 7.4, at room temperature for 10 min. Following three washes with ice-cold PBS, the cells were permeabilized in a permeabilization buffer (1x PBS, pH 7.4, 0.1% Triton X-100) at room temperature for 10 min. Following three more washes with PBS, the cells were incubated in 1% BSA in PBST (1x PBS, pH 7.4, 0.1% Tween® 20) at room temperature for 30 min. The cells were then incubated in primary antibody solution, FLAG (rabbit, 1:500, Thermo Scientific, Cat# PA1984B) and Rac1 (clone 23A8, mouse, 1:500, Sigma, Cat# 05389) in 1% BSA in PBST at 4°C overnight. After three washes with PBS, the cells were incubated in secondary antibody solutions, containing anti-mouse Alexa Fluor 647 (1:1000, Invitrogen, Cat# A-21235) and anti-rabbit Alexa Fluor 555 (1:500, Invitrogen, Cat# A27039) in PBST at 4°C overnight. Cells were washed three more times with PBS and stained with DAPI (ProLong™ Gold Antifade Mountant with DNA Stain DAPI, Thermo Fisher) for 10 min at room temperature. Cells were washed with PBS and analyzed using the confocal microscope (Zeiss LSM 780). To calculate Pearson’s correlation coefficients between FLAG-ARHGEF7 (green) and Rac1 (red), the images were first auto-thresholded (Intermodes) to isolate peaks of association from background before being run through the Just Another Colocalization Plugin (JACoP) in ImageJ software to analyze association between ARHGEF7 and Rac1.

### In vitro ARHGEF7 binding analysis

100 µM of purified ARHGEF7 DP or ARHGEF7 SDP was incubated with 1 mM oxidized glutathione (GSSG, adjusted to pH 7.4) in 1x PBS at room temperature for 1 h. The protein was dialyzed against 1x PBS at room temperature for 1 h to remove unbound GSSG. 100 µM of purified GST-Rac1 was incubated with 1 mM GDP, 1 mM GTPγS, or none at room temperature for 1 h. ARHGEF7 constructs and GST-Rac1 were incubated with Glutathione Sepharose™ 4 Fast Flow resin (Cytiva) at 4°C for 1 h. The resin was pelleted through centrifugation at 1,400 rpm for 3 min and then washed three times with PBS. Proteins were eluted in 50 µL SDS-loading dye (2x) containing 10 mM DTT. Eluted proteins were separated by SDS-PAGE and transferred to PVDF membrane for western blot analysis, using antibodies to FLAG (1:1000, Sigma, Cat# F1804) and Rac1 (1:1000, Cell Signaling, Cat# 4651S).

For binding analysis with PI(3,5)P_2_, dialyzed glutathionylated ARHGEF7 SDP (WT and C312S) was then incubated with biotinylated PI(3,5)P_2_ PolyPIPosomes (Echelon Biosciences, Cat# Y-P035) at room temperature for 30 min. The ARHGEF7 SDP was added to a solution containing high-capacity streptavidin agarose beads (Pierce™) and rotated at 4°C overnight. The beads were pelleted through centrifugation at 1,400 rpm for 3 min and then washed three times with PBS. To elute the proteins, the beads were resuspended in 50 µL SDS-loading dye (2x) containing 10 mM DTT and incubated at 95°C for 10 min. Eluted proteins were separated by SDS-PAGE.

### ARHGEF7 enzyme kinetic assay

To determine Michaelis-Menten kinetics, 100 µM of purified ARHGEF7 SDP was incubated without or with 1 mM oxidized glutathione (GSSG, pH 7.4) in 1x PBS at room temperature for 1 h. Proteins were dialyzed against 1x PBS at room temperature for 1 h, followed by the addition of 1 mM GTP. 400 µM of purified GST-Rac1 was incubated with 400 µM BODIPY™-GDP (Thermo Scientific) at room temperature for 1 h, followed by the addition of 1 mM MgCl_2_ to stop the loading of the GDP into GST-Rac1. All enzymatic reactions contained 1 mM GTP, 1 mM MgCl_2_, BODIPY-GDP-bound GST-Rac1, and ARHGEF7 SDP. The addition of the ARHGEF7 SDP (mixed with GTP) initiated the reaction. Reactions were performed in a 96-well plate, with fluorescence intensity measured (excitation – 488 nm, emission – 535 nm) using a Synergy H1 microplate reader (BioTek). First, BODIPY-GDP-bound GST-Rac1 was maintained at a constant 10 µM, with varying ARHGEF7 SDP WT concentrations (10 nM, 20 nM, and 50 nM) to ensure the reaction remained at its initial rate. Second, ARHGEF7 SDP was kept at a constant concentration of 10 nM, with varying Bodipy-GDP-bound GST-Rac1 concentrations (0-200 µM) to determine Michaelis-Menten kinetics. The slopes of the fluorescence intensity loss were used as reaction rates for ARHGEF7 SDP, which were plotted against the concentrations of Bodipy-GDP-bound GST-Rac1 in GraphPad Prism to perform nonlinear regression and determine the Michaelis-Menten parameters.

### Full-length ARHGEF7 purification and activity

MDA-MB-231 cells were transfected with ARHGEF7 WT (pcDNA3-FLAG-mARHGEF7), grown for 48 h, and lysed in Co-IP lysis buffer. Cell lysates from four 10-cm plates were combined and then incubated with FLAG primary antibody (1:200) at 4°C for 1 h. The solution was added to a slurry of protein G agarose beads (Pierce™) and rotated at 4°C overnight. The beads were pelleted through centrifugation at 1,400 rpm for 3 min and then washed three times with PBS. To elute the full-length ARHGEF7, the beads were resuspended in a solution of 1 mg/mL 3x FLAG-peptide (Apexbio, Cat# A60014) and 0.1% Tween® 20 and incubated at 4°C for 2 h. Following centrifugation at 1,400 rpm for 3 min, eluted proteins were collected and separated by SDS-PAGE and analyzed by western blot analysis, using FLAG antibody (Sigma, Cat# F1804) and actin antibody (1:1000). Full-length ARHGEF7 concentration was estimated through comparison to standards (purified FLAG-SMYD2). 100 nM of full-length ARHGEF7 was incubated without and with 1 mM oxidized glutathione (GSSG, pH 7.4) in 1x PBS at room temperature for 1 h to induce S-glutathionylation. The enzymatic activity was measured, as described above, in reactions containing 1 mM GTP, 1 mM MgCl_2_, 200 µM Bodipy-GDP-bound GST-Rac1, and 10 nM full-length ARHGEF7.

### Statistical analysis

All data are presented as means ± SD and were statistically analyzed using one-way ANOVA or two-way ANOVA followed by Tukey’s *post-hoc* test. The value p < 0.03 is statistically significant.

## Data availability

All data supporting the findings of this study are available from the corresponding author upon request.

## Supplementary information

This article contains supporting information.

## Acknowledgements

The research was supported by the National Institutes of Health (NIH) (R01 GM143214) (Y.H.A) and a research fund from Drexel University (Y.H.A). We would like to thank all members of the Ahn group for their assistance with the experiments.

## Author Contributions

Conceptualization, W.S., Y.H.A.; Formal Analysis, W.S., M.C.S., F.M.R., D.S.K.K., R.P., Y.H.A.; investigation, W.S., M.C.S., F.M.R., D.S.K.K., R.P.; supervision, Y.H.A.; visualization, W.S., M.C.S., Y.H.A.; writing – original draft, W.S.; writing – review & editing, W.S., M.C.S., F.M.R., D.S.K.K., R.P., Y.H.A.; funding acquisition, Y.H.A.

