## Supplementary Information for "ARHGEF7 S-glutathionylation promotes cancer cell migration through Rac1 activation"

Schiff *et al*.

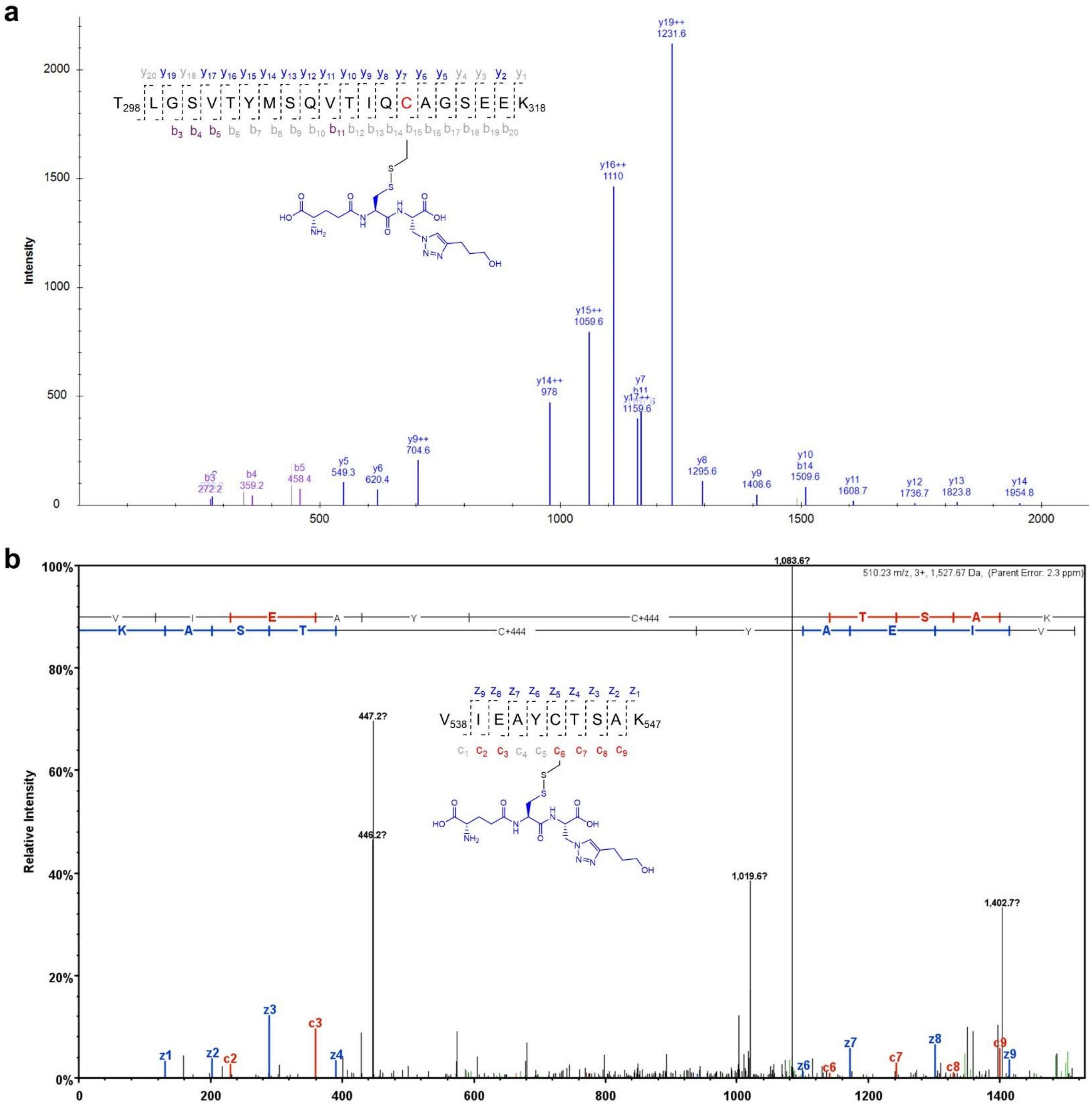


**Supplementary Figure 1. Tandem MS/MS spectrum of glutathionylated peptides in ARHGEF7.** (**a**) Tandem mass analysis of ARHGEF7 C312 glutathionylation (**b**) Tandem mass analysis of ARHGEF7 C543 glutathionylation. The data was retrieved from our previous proteomic data (C312[S1] and C543[S2]). Cysteine numbering is based on human isoform-a or mouse isoform-c (total 646 residues). Mouse ARHGEF7 glutathionylated peptides were detected in the mouse HL-1 cell line.[S1, 2]

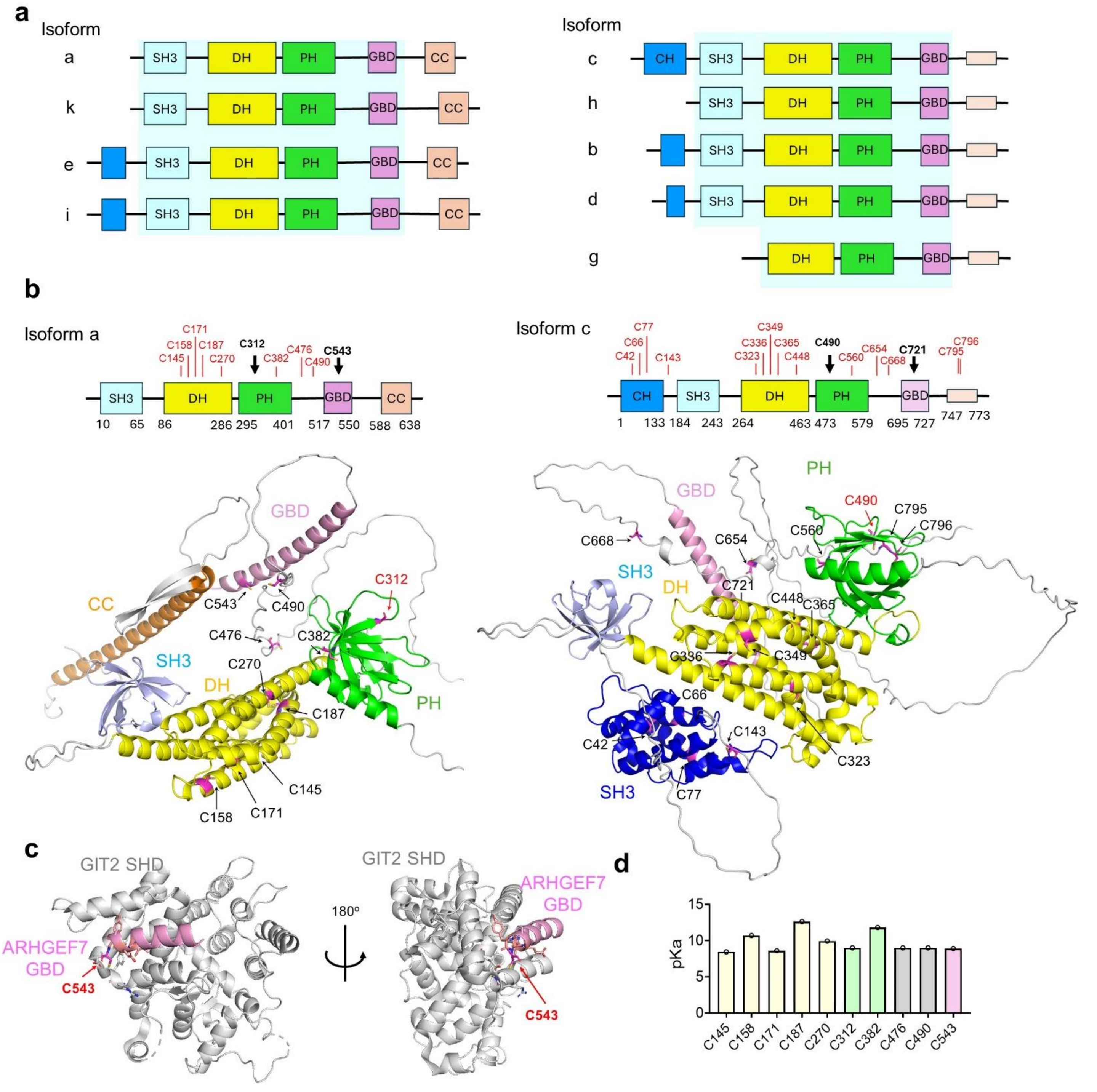


**Supplemental Figure 2. ARHGEF7 isoform, structure, and its cysteine $pK_a$ analysis.** (**a**) Different isoforms of human ARHGEF7. Isoform sequences were retrieved from NCBI gene databases and compared to note the differences in domains and flexible regions. SH3-DH-PH-GBD are identical, except for isoform-g. Similar sequences in N-terminus (before SH3) or C-terminus (after GBD) were noted among a, k, e, and I (left) or c, h, b, d, and g (right). (**b**) Domain and structure of isoform-a and -c. Cysteine sites were indicated by arrows. Cysteines identified for glutathionylation in mass spectrometry were indicated by black arrows. Structures were retrieved from the Alpha-Fold database. (**c**) The structure of the GIT1-ARHGEF7 complex. The GIT SHD domain structure is shown with GBD of ARHGEF7 containing C543 (PDB: 6JMT).[S3] (**d**) ARHGEF7 cysteine $pK_a$ analysis. ARHGEF7 Alpha-Fold structure was used to predict the $pK_a$ values of cysteines using the ProPKa program.[S4]

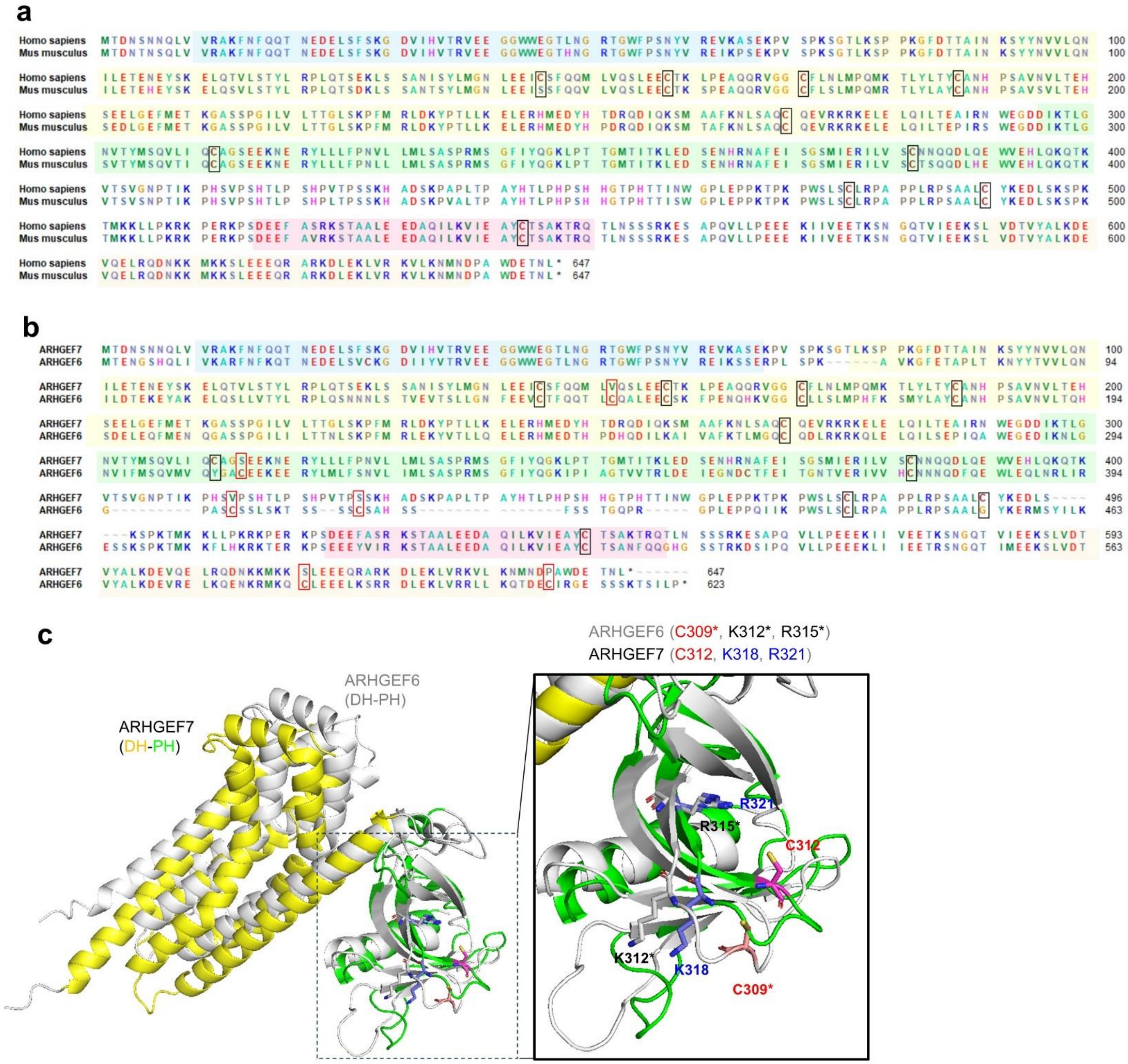


**Supplemental Figure 3. ARHGEF7 amino acid sequence with its orthologs and its family.** (**a**) The amino acid sequence comparison between human (isoform-a, 646 residues, NM_001113513) and mouse (isoform-c, 646 residues, NM_017402) ARHGEF7 orthologs. Cysteines are marked with boxes. (**b**) The amino acid sequences between human ARHGEF7 (βPix, NM_001113513) and human ARHGEF6 (αPix, NM_001306177). ARHGEF7 C312 is not aligned, but close to ARHGEF6 C309. (**c**) Structural alignment of ARHGEF7 DH-PH (yellow-green, AF-Q14155-1-F1-model_v6) and ARHGEF6 DH-PH domains (gray, AF-Q15052-2-F1-model_v6). ARHGEF7 C312 and ARHGEF6 C309 are closely aligned in structures with a similar arrangement of neighboring basic residues (K and R) that may interact with $PIP_2$. Structures were retrieved from the Alpha-Fold database.

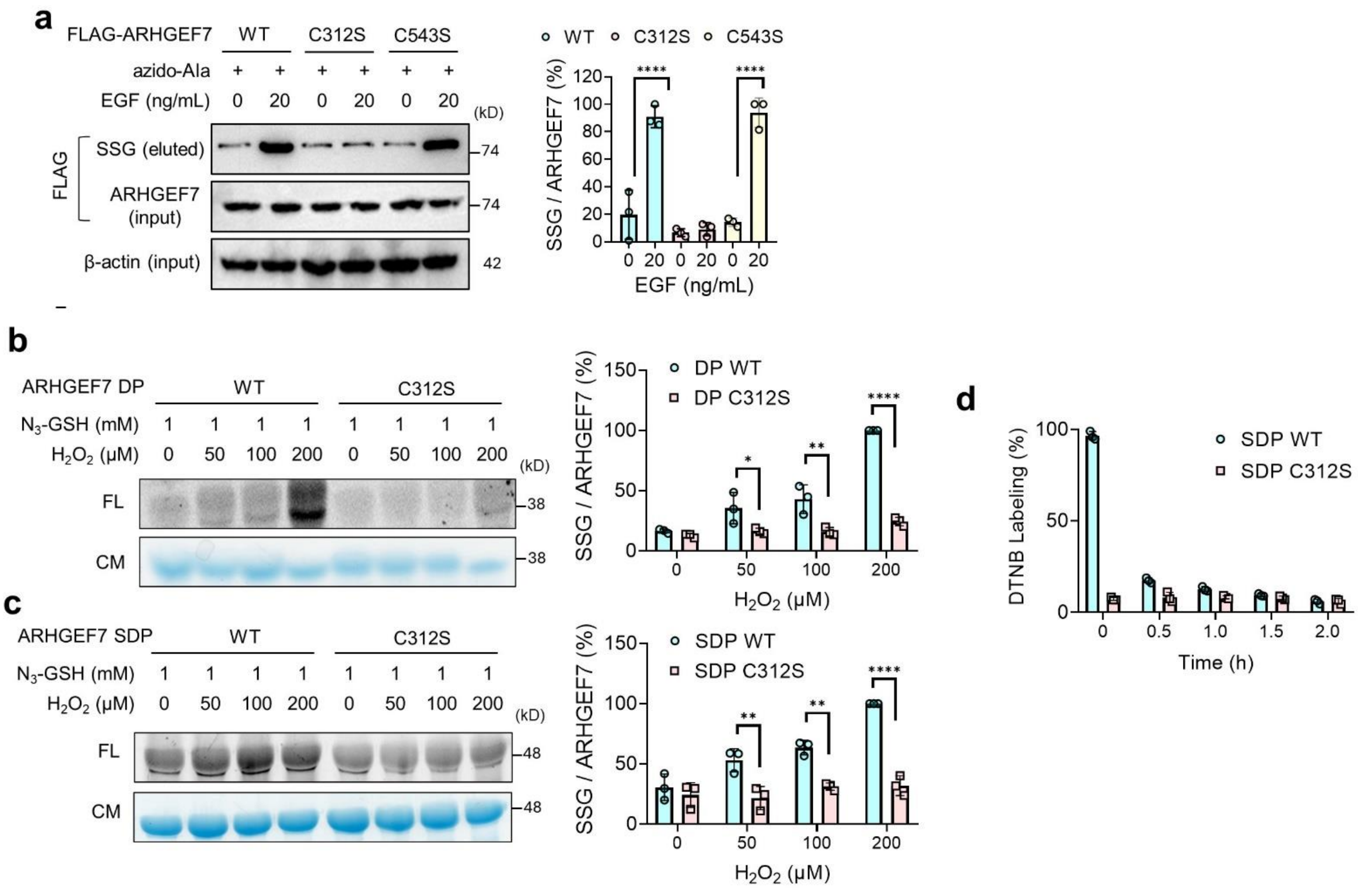


**Supplemental Figure 4. ARHGEF7 S-glutathionylation analysis.** (**a**) ARHGEF7 S-glutathionylation in response to a short incubation time of EGF. MDA-MB-231 cells expressing WT, C312S, or C543S were treated with EGF for 30 min (n=3). Lysates were subjected to click reactions with biotin-alkyne. Glutathionylated proteins were analyzed before (input) and after (eluted) streptavidin-enrichment. (**b**-**c**) S-glutathionylation analysis of purified ARHGEF7 constructs in vitro. ARHGEF7 DP (**b**) or SDP (**c**) in the presence of azido-glutathione (1 mM) was incubated with hydrogen peroxide ($H_2O_2$). After click reactions with rhodamine-alkyne, glutathionylated proteins were analyzed by fluorescence (FL) and Coomassie stain (CM). (**d**) Free sulfhydryl level analysis of ARHGEF7 constructs upon incubation of oxidized glutathione (GSSG). Purified ARHGEF7 SDP WT and C312S were individually incubated with GSSG (1 mM) for the indicated times, followed by a reaction with Ellman's reagent (DTNB) to monitor the levels of remaining reduced cysteines. The DTNB reaction product was detected through absorbance at 412 nm. The statistical difference was analyzed by one-way ANOVA (**a**) or two-way ANOVA (**b-d**) with Tukey's post-hoc test , where $*p < 0.03$, $**p < 0.002$, $***p < 0.0002$, $****p < 0.0001$.

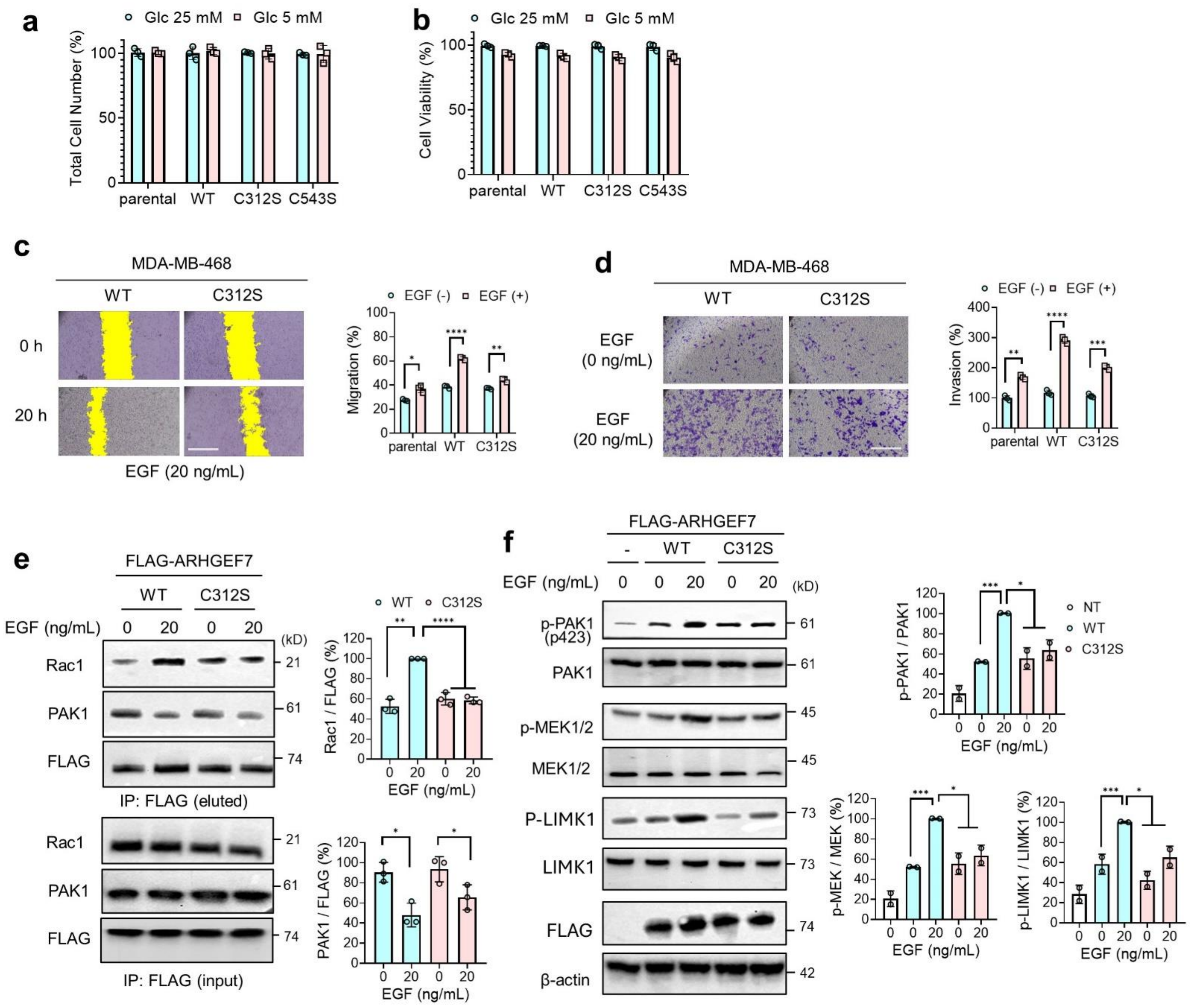


**Supplemental Figure 5. Cell phenotype and mechanistic analysis after ARHGEF7 glutathionylation.** (**a-b**) Cell number and viability analysis. MDA-MB-231 cells expressing ARHGEF7 constructs were incubated in high glucose (25 mM, HG) or low glucose (5 mM, LG) conditions for 24 h. Total cell number (n=3) (**a**) and cell viability by trypan blue (n=3) (**b**) were measured. (**c-d**) Migration and Invasion analysis of MDA-MB-468 upon ARHGEF7 C312 glutathionylation. MDA-MB-468 cells expressing the ARHGEF7 construct were analyzed for migration by an in vitro scratch assay (n=3) (**c**) and for invasion by a transwell invasion assay (n=3) (**d**). Yellow colors indicate the area without cells. A scale bar = 500 μm. (**e-f**) Mechanistic Rac1-PAK1 signaling pathway analysis in response to EGF. MDA-MB-231 cells expressing ARHGEF7 WT or C312S were incubated with EGF for 20 h. (**e**) ARHGEF7 co-immunoprecipitation (co-IP) with Rac1 and PAK1 (n=3). (**f**) Phosphorylation levels of PAK1, LIMK1, and MEK1/2 were analyzed by western blot (n=2). The statistical difference was analyzed by two-way ANOVA (**a-d**) or one-way ANOVA (**e-f**) with Tukey's post-hoc test, where *$p < 0.03$, **$p < 0.002$, ***$p < 0.0002$, ****$p < 0.0001$.

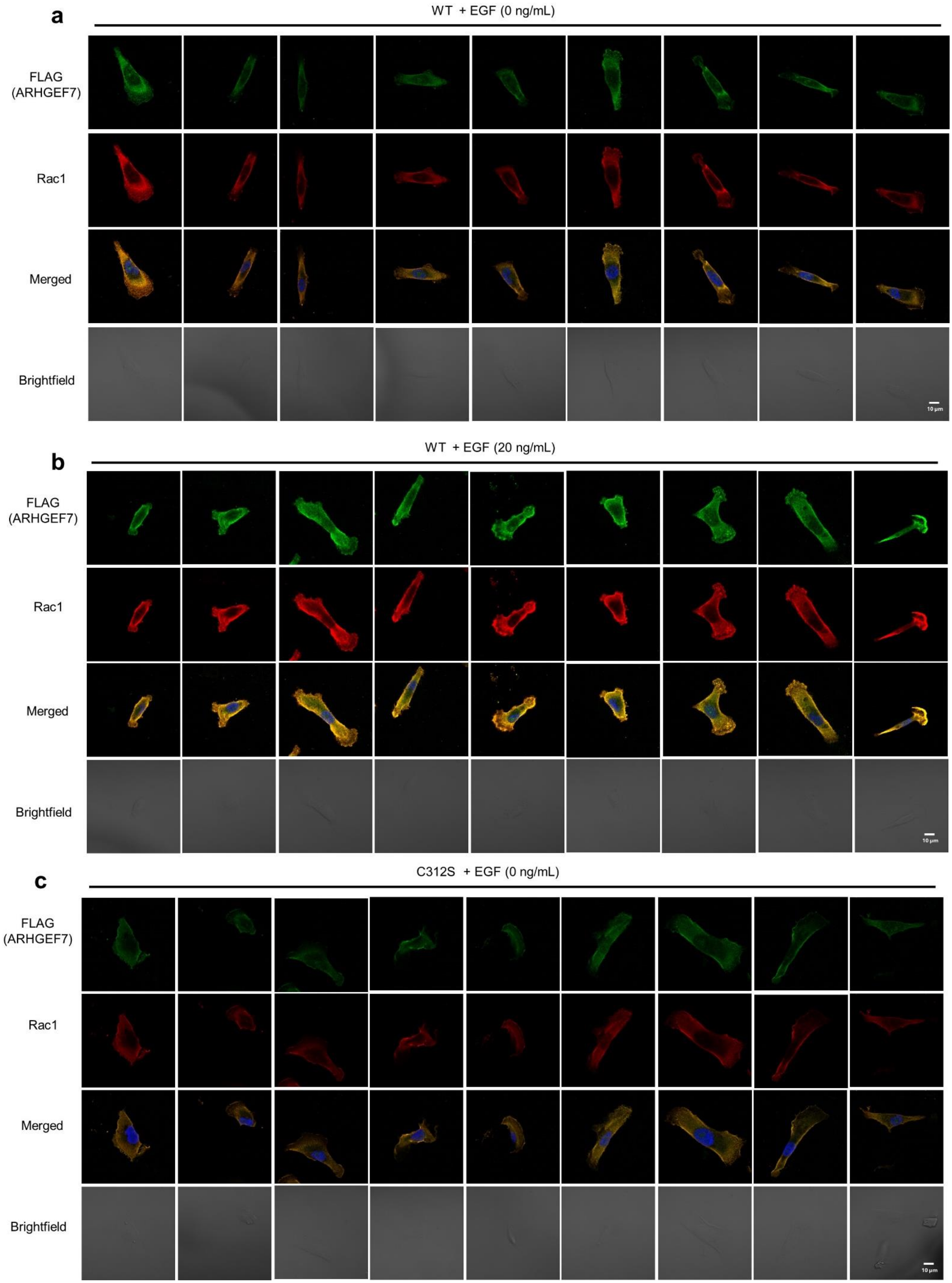
a
WT + EGF (0 ng/mL)
FLAG (ARHGEF7)
Rac1
Merged
Brightfield
10 µm
b
WT + EGF (20 ng/mL)
FLAG (ARHGEF7)
Rac1
Merged
Brightfield
10 µm
c
C312S + EGF (0 ng/mL)
FLAG (ARHGEF7)
Rac1
Merged
Brightfield
10 µm

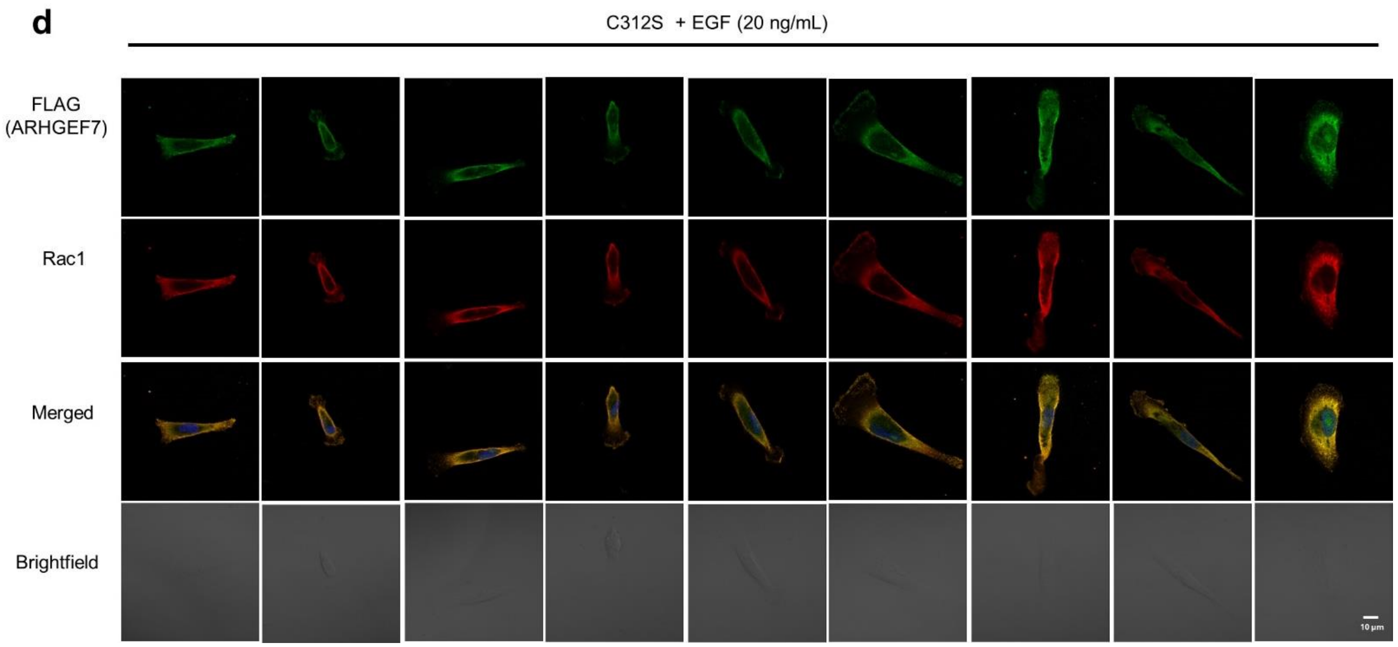


**Supplemental Figure 6. Localization analysis of ARHGEF7 and Rac1.** MDA-MB-231 cells expressing FLAG-ARHGEF7 WT (**a**-**b**) or C312S (**c**-**d**) were incubated with none or EGF (20 ng/mL) for 16 h. Cells were fixed and analyzed by immunostaining with antibodies to FLAG (Alexa Fluor 555, green) and Rac1 (Alexa Fluor 647, red). DAPI (blue) was added to visualize the nucleus. Replicate images are shown and include the data in Figure 3D. The representative images and quantification analysis are shown in Figure 3D. A scale bar = 10 µm.

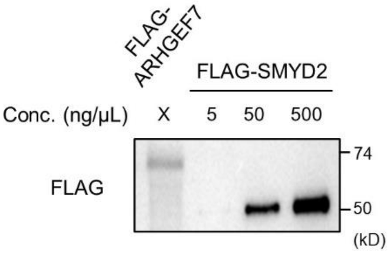


**Supplemental Figure 7. Analysis of purified ARHGEF7 construct.** Validation of FLAG-ARHGEF7 purification. FLAG-ARHGEF7 overexpressed in MDA-MB-231 cells was purified by FLAG antibody-based affinity purification and eluted by 3xFLAG peptide. The eluate was analyzed by Western blot with FLAG antibody. The eluted FLAG-ARHGEF7 concentration was estimated by comparing to a positive control of purified FLAG-SMYD2 in a series of concentrations (5, 50, 500 ng/µL) (all loaded in equal volumes), estimating approximately 10 ng/µL.

### Supplementary Methods

**Free thiol assay.** 100 µg of ARHGEF7 DP or SDP was incubated with 1 mM oxidized glutathione (GSSG, adjusted to pH 7.4) in 1x PBS at room temperature for various times up to 2 h. The protein was dialyzed against 1x PBS at room temperature for 1 h to remove unbound GSSG. 2 mM Ellman's Reagent (DTNB) was dissolved in a DNTB reaction buffer (100 mM Tris, pH 8.0, 50 mM sodium acetate) before mixing with ARHGEF7 DP or SDP for 10 min. Absorbance was measured at 412 nm to quantify DTNB-modified cysteines in ARHGEF7 using a Synergy H1 microplate reader (BioTek).

**Bioinformatic analysis.** pKa values of cysteines were analyzed by the PROPKa program (https://www.ddl.unimi.it/vegaol/propka.htm).[S4] Accessible surface area (ASA) values of cysteines were analyzed by the Adaptive Poisson-Boltzmann Solver (APBS) software (https://server.poissonboltzmann.org/).[S5] An AlphaFold structural model (AF-Q14155-1-F-model_v6_isoform_a) was used to calculate the $pK_a$ and ASA values.
